# “Characterisation of the biflavonoid hinokiflavone as a pre-mRNA splicing modulator that inhibits SENP”

**DOI:** 10.1101/123026

**Authors:** Andrea Pawellek, Ursula Ryder, Triin Tammsalu, Lewis J. King, Helmi Kreinin, Tony Ly, Ronald T. Hay, Richard Hartley, Angus I. Lamond

**Affiliations:** University of Dundee, School of Life Sciences, Centre for Gene Regulation and Expression, Dundee, DD1 5EH, UK; University of Glasgow, WestCHEM School of Chemistry, Glasgow, G12 8QQ, UK

## Abstract

Here, we identify the plant biflavanoid hinokiflavone as an inhibitor of splicing *in vitro* and modulater of alternative splicing in multiple human cell lines. Hinokiflavone inhibits splicing *in vitro* by blocking one or more early steps of spliceosome assembly, leading to accumulation of the A complex. Multiple human cell lines treated with hinokiflavone show changes in the alternative splicing of different pre-mRNA substrates, but little or no change in transcription. They also show altered subnuclear organization, specifically of splicing factors required for A complex formation, which relocalized together with SUMO1 and SUMO2 into enlarged nuclear speckles. While most cell lines treated with hinokiflavone showed cell cycle arrest and eventual cell death, dependent on time and concentration, the promyelocytic NB4 cell line, which expresses the SUMO target PML-RARalpha fusion protein, was exquisitely sensitive to apoptosis following hinokiflavone treatment. Hinokiflavone treatment increased protein SUMOylation levels, both in *in vitro* splicing reactions and in cells, with little or no effect on levels of ubiquitinylated proteins. Hinokiflavone also inhibited the catalytic activity of purified *E. coli* expressed SUMO protease, SENP1 *in vitro*, indicating the increase in SUMOylated proteins results primarily from inhibition of de-SUMOylation. Using a quantitative proteomics assay we identified many SUMO2 sites whose levels increased following hinokiflavone treatment, with the major targets including 6 proteins that are associated with U2 snRNP and required for A complex formation. These data identify hinokiflavone as a SUMO protease inhibitor and indicate SUMOylation of splicing factors may be important for modulating splice site selection.

## Introduction

Pre-mRNA splicing is an essential step in gene expression in eukaryotes. During splicing, intron sequences are removed from nascent, pre-mRNA gene transcripts via two, sequential transesterification reactions, thereby joining exon sequences to generate messenger RNA (mRNA) for protein translation (reviewed in ^1,2,3^). The splicing of pre-mRNA takes place in the cell nucleus and is catalyzed by the large (>3 MDa), ribonucleoprotein spliceosome complex. A separate spliceosome complex independently forms on each intron in a pre-mRNA transcript, involving a stepwise assembly pathway of the U1, U2 and U4/5/6 snRNP spliceosome subunits, together with additional protein splicing factors. The core splicing machinery, spliceosome assembly pathway and reaction mechanism is highly conserved across eukaryotes.

In higher eukaryotes, most pre-mRNA transcripts have multiple introns and it is common for a single gene to give rise to multiple different mRNA products through alternative patterns of intron removal, e.g. via either the choice of different 5’ and/or 3’ splice sites, or via exon skipping etc. (reviewed in ^3,4^). Alternative splicing is thus a major mechanism for generating proteome diversity in higher eukaryotes and is likely to have played a major role in the evolution of organismal complexity. Changes in the pattern of alternative splicing of pre-mRNAs is highly regulated and plays an important role during organismal development, cellular differentiation and the response to many physiological stimuli. For example, a number of genes regulating apoptosis pathways, such as Bcl2, can be alternatively spliced to generate distinct mRNAs encoding proteins that either promote, or inhibit, cell survival (^5,6^).

Recent high-throughput deep sequencing experiments have shown that almost all human genes produce alternatively spliced mRNA isoforms^7^. The dysregulation of alternative splicing is now recognized as a major mechanism for multiple forms of human disease, including inherited disorders, cancer, diabetes and neurodegenerative diseases^8^. The ability to target the modulation of specific classes of alternative pre-mRNA splicing events using small molecules is thus seen as an important new therapeutic strategy for drug development and future disease therapy (^9,10,11^).

Chemical compounds are widely used as biotools to study the gene expression machineries involved in transcription and translation. In contrast, well-characterized compounds that can be used to dissect and study the splicing machinery are still very limited. Most splicing inhibitors described to date are natural products. The best-studied group are all SF3B1 inhibitors, which have either been isolated from the broth of bacteria, or are their synthetic derivatives, like Spliceostatin A, E7107 and herboxidine (^12,13,14^). In addition to the SF3B1 inhibitors, another plant derived compound, the biflavone isoginkgetin (extracted from the leaves of the ginko tree), has been reported to inhibit splicing, but its target remains unknown^2^.

Biflavones such as isoginkgetin belong to a subclass of the plant flavonoid family, which have been reported to possess a broad spectrum of pharmacological activity, including antibacterial, anticancer, antiviral and anti-inflammatory functions^15^. In view of their broad biological activity, we tested a set of biflavones for their ability to change pre-mRNA splicing *in vitro* and *in cellulo*. This identified the biflavone hinokiflavone as a new general splicing modulator active both *in vitro* and *in cellulo*. Further analysis indicated that hinokiflavone promotes unique changes in subnuclear structure and dramatically increases SUMOylation of a subset of spliceosome proteins by inhibiting SENP protease activity.

## Results

### Biflavonoids as splicing modulators

We tested a set of bioflavonoids for a potential effect on pre-mRNA splicing *in vitro*, including amentoflavone, cupressuflavone, hinokiflavone and sciadopitysin (Figure 1A). As a positive control we included the previously reported biflavonoid splicing inhibitor, isoginkgetin^16^. Initially, we screened each compound at a high final concentration (500 μM), for a potential inhibitory effect on splicing of the model Ad1 and HPV18 E6 pre-mRNAs, using HeLa nuclear extract in conjunction with a non-radioactive RT-PCR *in vitro* splicing assay (see Methods). Cupressuflavone did not inhibit splicing under these conditions, while hinokiflavone, amentoflavone, isoginkgetin and sciadopitysin all showed some inhibition of *in vitro* splicing of the adenovirus and/or the HPV18E6 pre-mRNAs. The strongest inhibitory effects were obtained with hinokiflavone and amentoflavone, which had not previously been identified as affecting the spliceosome (Figure 1B). We note that the active compounds each have a linkage connecting the B and A’ units of the two flavone moieties.

**Figure 1.**
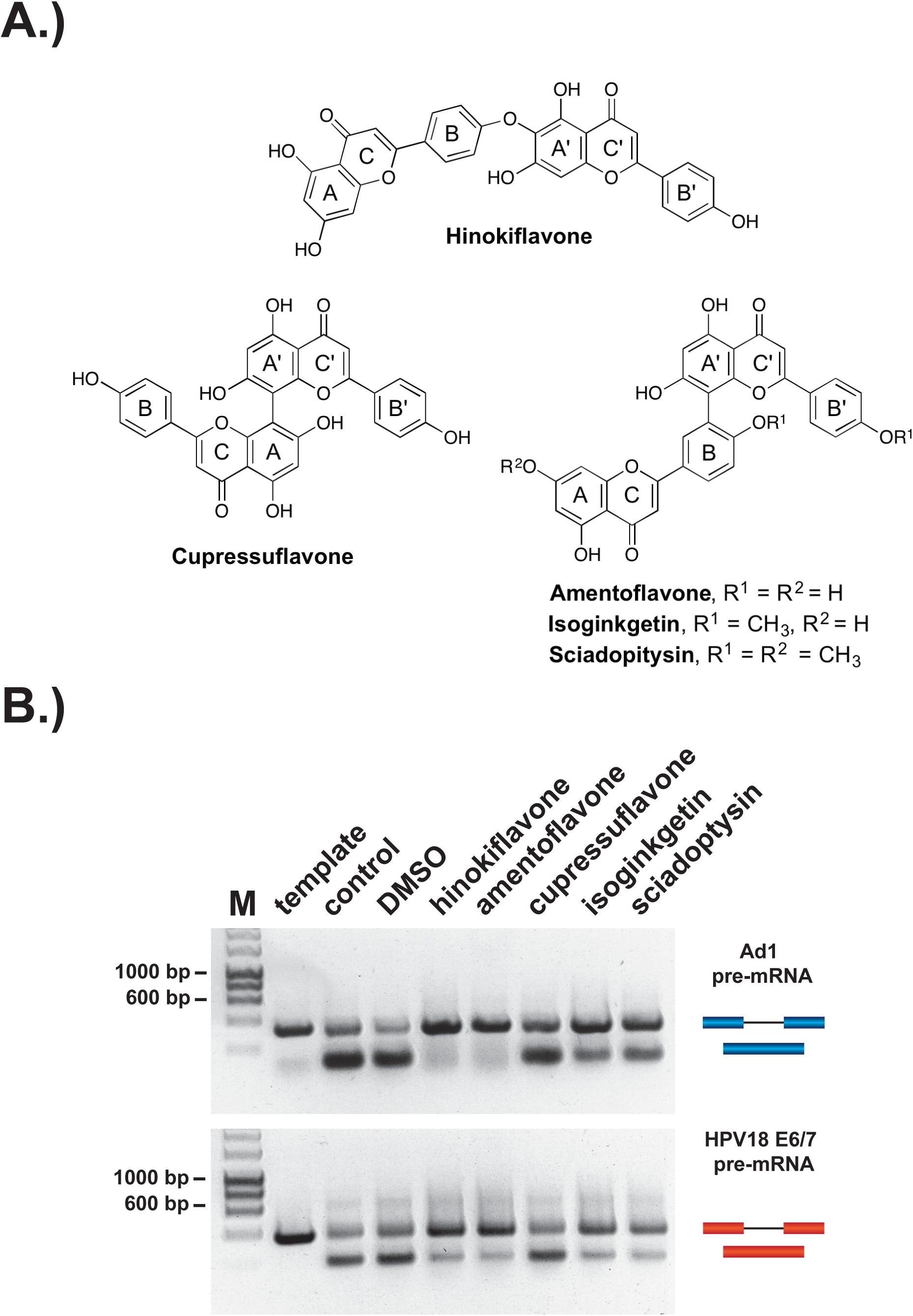
Biflavones inhibit splicing in vitro. (A) Chemical structures of the 5 biflavones, hinokiflavone, amentoflavone, cupressuflavone, isoginkgetin and sciadopitysin were tested for their ability to inhibit splicing of the Ad1 and the HPV18 E6 pre-mRNAs in a non radioactive RT-PCR based *in vitro splicing* assay. (B) Splicing assays show that all compounds except cupressuflavone inhibited splicing to varying degrees in this assay system.

Next, we tested the biflavonoids for their ability to alter pre-mRNA splicing in human cell lines. First, HEK293 cells were treated for 24h with 20-100 μΜ of each compound and a RT-PCR assay was used to detect potential changes in the splicing patterns of HSP40, RIOK3, RBM5, FAS and MCL1 pre-mRNAs. With the set of pre-mRNAs tested, only isoginkgetin and hinokiflavone induced changes in alternative splicing patterns *in cellulo*, whereas amentoflavone, cupressuflavone and sciadopitysin did not alter splicing of the tested transcripts (Supplementary Figure1).

As hinokiflavone was the only compound to show strong effects on pre-mRNA splicing both *in vitro* and *in cellulo*, we concentrated on characterizing its effect on modulating splicing in more detail. First, HeLa, HEK293 and NB4 cells were treated with either a DMSO control, or with hinokiflavone at either 10 μM, 20 μM or 30 μM for 24h, then cells harvested and total RNA extracted. A semiquantitative RT-PCR assay was performed, using primer pairs to detect either changes in alternative pre-mRNA splicing for transcripts from the MCL1, NOP56, EIF4A2 and FAS genes, or intron retention for transcripts from the HSP40, RIOK3, ACTR1b and DXO genes. Hinokiflavone modulated splicing in all three cell lines, but with the greatest effect observed in NB4 cells (Figure 2).

**Figure 2.**
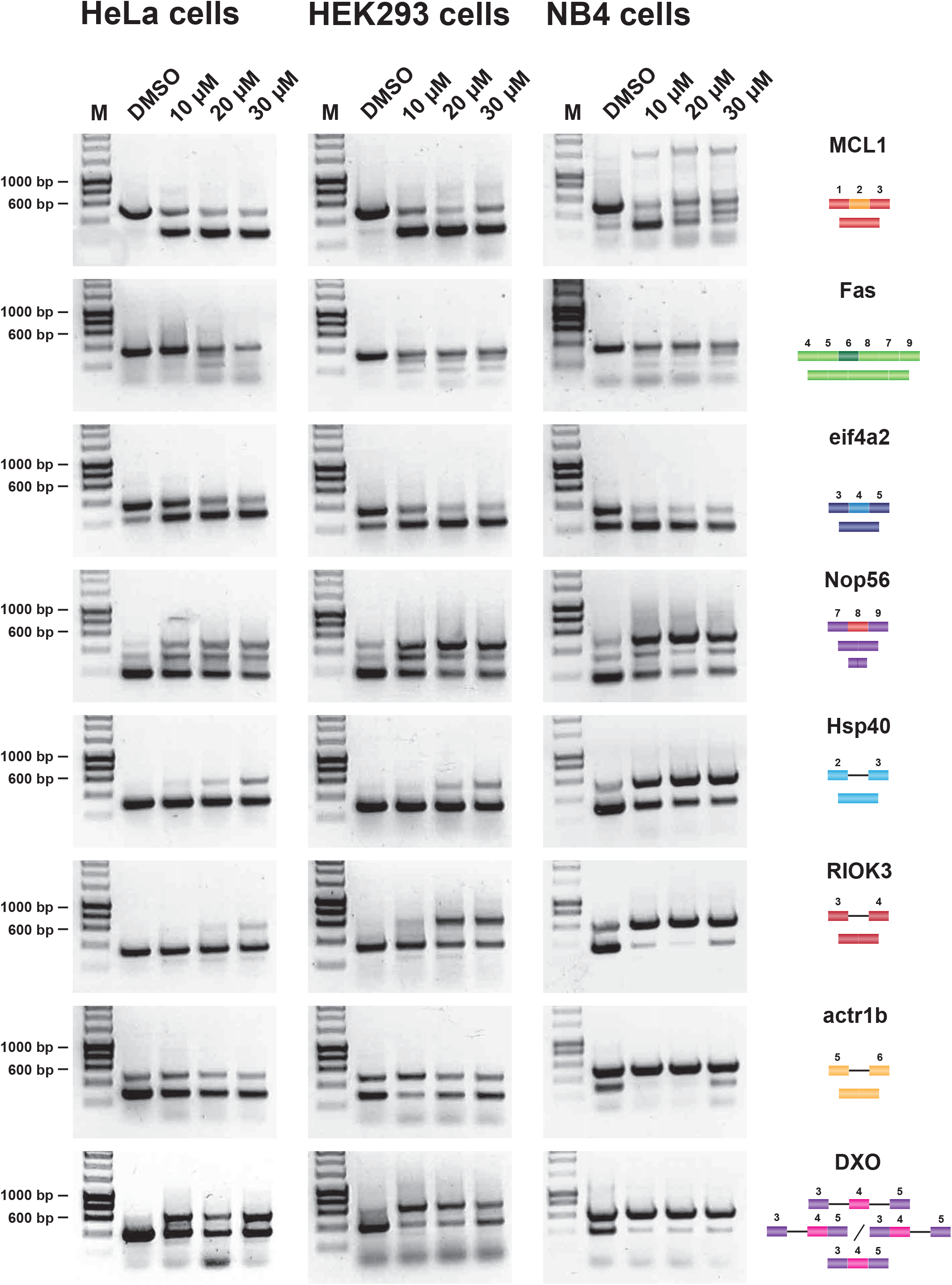
Hinokiflavone modulates splicing in cells. Semiquantitative RT-PCR analysis of cells treated with increasing concentrations of either hinokiflavone, or DMSO control, for 24h. HeLa cells and HEK293 cells were examined for intron inclusion of HSP40, RIOK3, ACTR1b and DOX pre-mRNAs and for exon skipping of MCL1, NOP56, EIF4A2 and RBM5 pre-mRNAs. The position of different cDNA products are pictured on the right of each gel image, and the molecular weight markers are shown on the left. Lane M, marker (hyperladder, 1kb)

An interesting differential effect on the alternative splicing of MCL1 was observed between the different cell lines. Thus, in HeLa and HEK293 cells, hinokiflavone promoted exon 2 skipping, whereas, in NB4 cells, multiple, aberrant alternatively spliced MCL1 isoforms were detected. In the presence of 10 μM hinokiflavone a third band of around 1.5kb was detected next to the expected 2 MCL1 isoforms. In addition to the 1.5kb band, a loss of the MCL1-S isoform and the appearance of a larger splice variant migrating between MCL1-S and MCL1-L was observed in cells treated with either 20 μM, or 30 μM hinokiflavone.

Intron retention in the Hsp40 and RIOK3 transcripts was mainly detected in HeLa cells at 30 μM, in HEK293 cells at both 20 μM and 30 μM and in NB4 cells at all concentrations of hinokiflavone tested. In the case of ACTR1B, no changes in alternative splicing were detected in HeLa cells, whereas in HEK293 cells, intron inclusion was only observed in the presence of 10 μM hinokiflavone, while in NB4 cells intron inclusion was observed at all concentrations tested. Hinokiflavone also promoted intron 3 and intron 4 retention in the DXO transcripts and exon 4 skipping in the EIF4A2 transcripts in all cell lines and at all concentration tested. In the case of NOP56, hinokiflavone not only induced an increase in the production of NOP56 transcripts with retained exon 8, but also induced the usage of alternative 5’ and 3’ splice sites. In the case of FAS, multiple PCR products were detected, which were individually cloned and sequenced (Supplementary Figure 2). The FAS isoforms identified were either lacking exon 5, 6, 7 or 8, exon 6 and exon 7, or exons 6, 7 and 8.

To ensure that these observed effects on splicing *in vitro* and *in cellulo* were indeed caused by hinokiflavone, rather than by some minor product in the commercially available hinokiflavone isolated from a natural source, we developed a synthetic route for generating the hinokiflavone molecule. A detailed description of the synthetic route will be published separately (King et al., manuscript in preparation). Importantly, we find that chemically synthesized hinokiflavone is spectroscopically identical to hinokiflavone isolated from a natural source and has the same effects on pre-mRNA splicing in cells (Supplementary Figure 3). We conclude that hinokiflavone is able to modulate pre-mRNA splicing activity.

### Hinokiflavone stalls spliceosome assembly at the A complex

To investigate whether hinokiflavone inhibits splicing by preventing spliceosome assembly, *in vitro* splicing reactions were carried out using radioactive Ad1 pre-mRNA and either DMSO (control), or 500 μΜ hinokiflavone, then analyzed both by denaturing PAGE to detect reaction products and by native gel electrophoresis to monitor spliceosome assembly (Figure 3). Hinokiflavone inhibited the formation of both splicing products and intermediates, with no inhibition seen with the DMSO control, in comparison with untreated nuclear extract (Figure 3A). After 1h incubation, analysis using native gels showed the typical pattern of A, B and C spliceosome complexes in the DMSO control, similar to untreated nuclear extract. However, in the hinokiflavone treated extract, only H/E and A complexes were detected (Figure 3B). This indicates that the inhibition of splicing caused by hinokiflavone results from a failure in the transition from the A to the B complex during spliceosome assembly.

**Figure 3.**
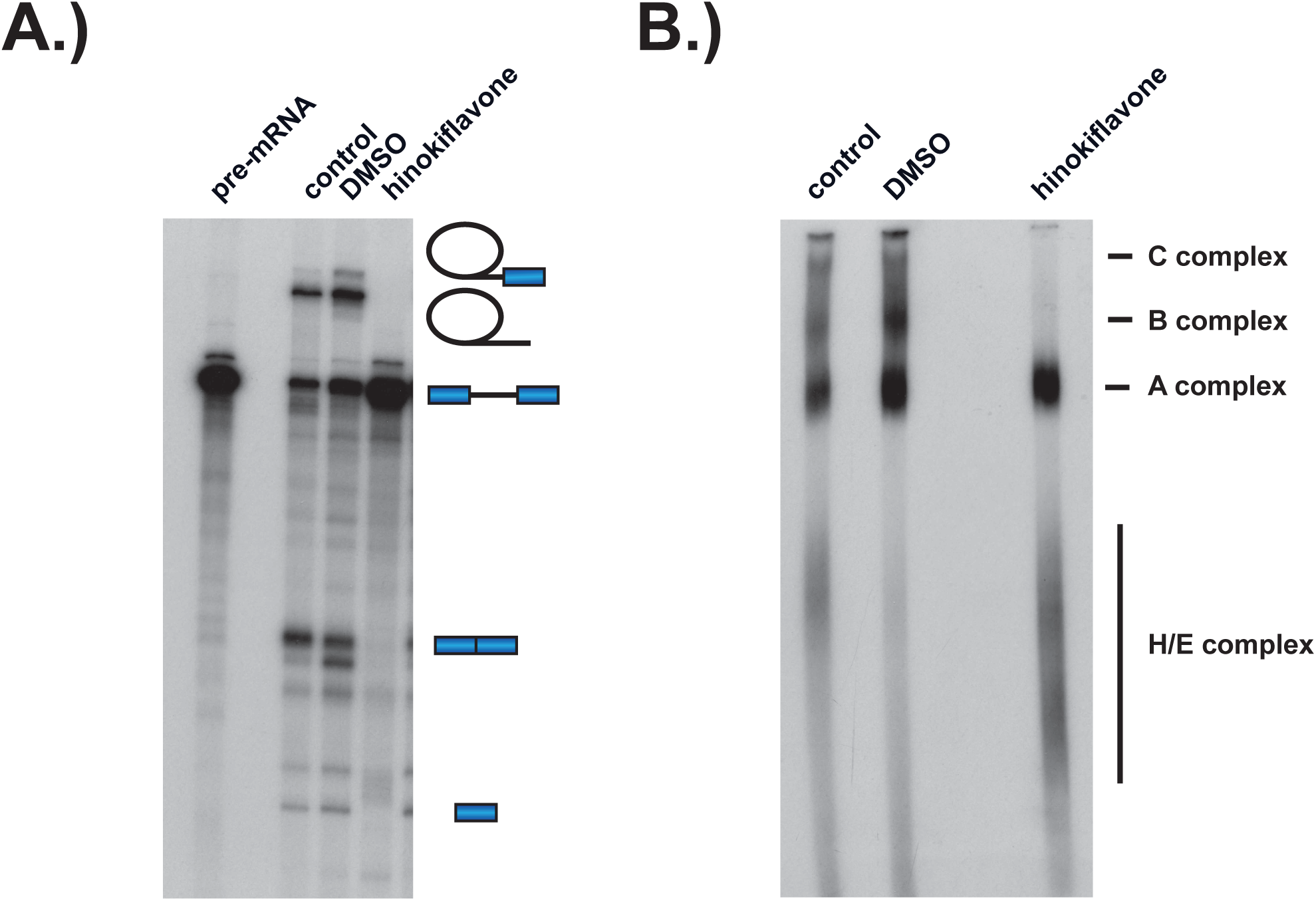
Hinokiflavone stalls spliceosome assembly after A complex formation. Formation of splicing complexes on the Ad1 pre-mRNA was analysed on a native agarose gel after incubation with either DMSO (control) or 500 μM hinokiflavone. The positions of the splicing complexes C, B, A and H/E are indicated on the right.

### Hinokiflavone blocks cell cycle progression

Next, we tested the effect of hinokiflavone on cell cycle progression. HeLa, HEK293 and NB4 cells were each treated either with DMSO as a negative control, or with hinokiflavone, at a final concentration of 10 μM, for 24h. The cells were then fixed, labelled with propidium iodide and analyzed by flow cytometry (Figure 4). Interestingly, hinokiflavone differentially affected the cell lines tested, with most showing either cell cycle arrest, or eventual cell death, dependent upon concentration and time of treatment. The most dramatic effect, however, was observed for the acute promyelocytic cell line NB4, where most cells become apoptotic after 24h exposure to 10 μM hinokiflavone (Figure 4).

**Figure 4.**
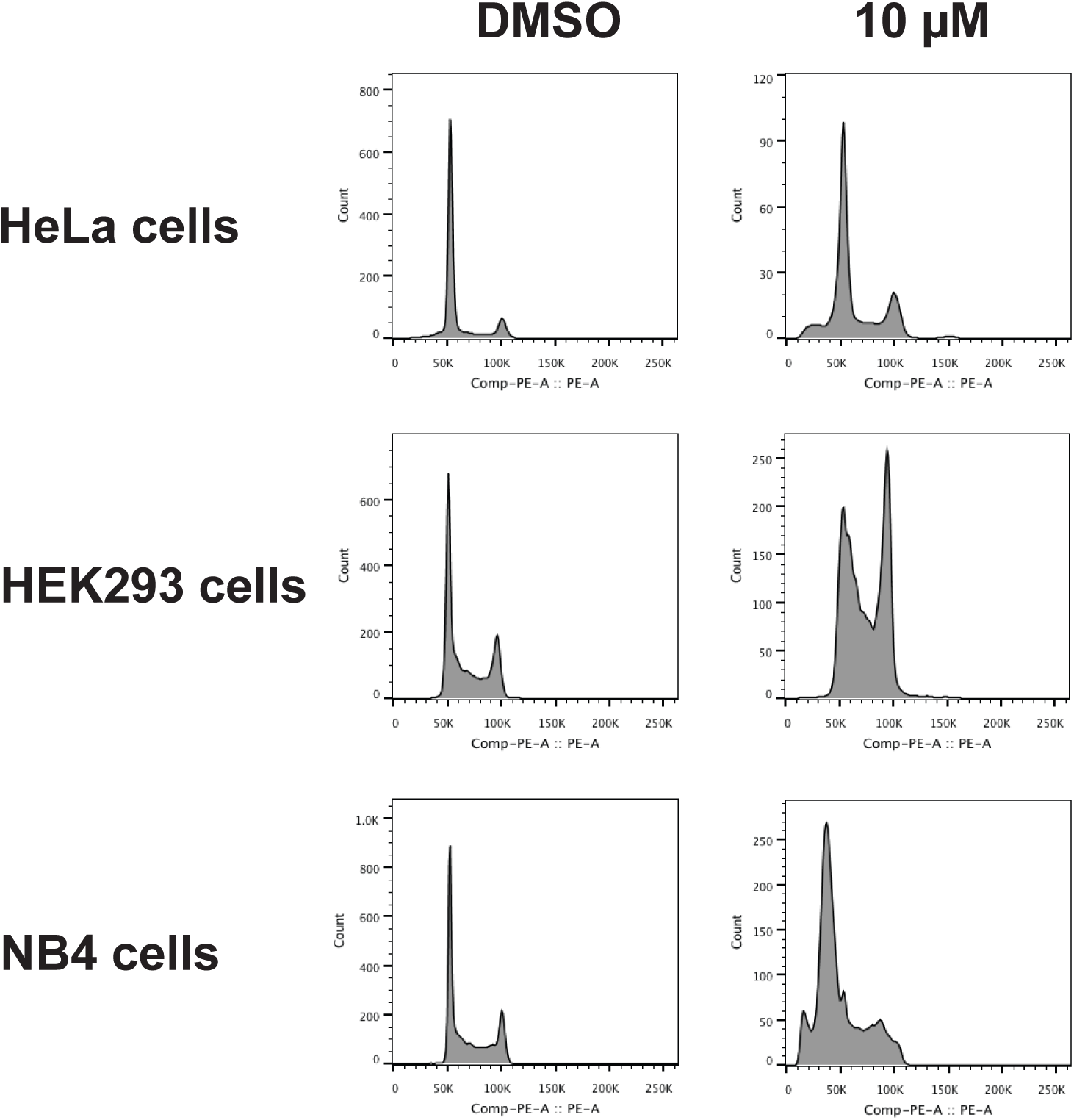
Hinokiflavone shows cell cycle specific effects. Cell cycle analysis was performed on HeLa, HEK293 and NB4 cells treated with either 10 μM hinokiflavone, or DMSO (control), for 24h. Cellular DNA content was measured by propodium iodide staining followed by flow cytometry analysis.

### Hinokiflavone alters nuclear organization of a subset of splicing factors

We examined the effect of hinokiflavone treatment on subcellular organization, in particular, the subnuclear organization of splicing factors and other nuclear components. For this, HeLa cells were treated with 20 μM hinokiflavone for 24h then the cells were fixed, permeabilised and stained with antibodies specific for the splicing factors SRSF2 (SC35), DDX46, U2AF65, SART1, SR proteins, CDC5L, PLRG1, BCAS2, PRP19, CTNNBL1 and snRNP200 (Figure 5). This showed a change in the ‘speckled’ nuclear staining pattern typical of many splicing factors, with the formation of enlarged and rounded ‘mega speckles’ (Fig 5A). Interestingly, two groups of splicing factors could be distinguished, based upon their response to hinokiflavone. The first group, including SRSF2 (SC35), U1A, DDX46, U2AF65 (U2AF2) and SR proteins, which are all involved in early steps of spliceosome assembly, were relocalized into the mega speckles. However, a second group, including CDC5L, PLRG1, BCAS2, PRP19, CTNNBL1 and snRNP200, which are all associated with later steps of the splicing process and assemble into the spliceosome after A complex formation, were not enriched in the mega speckles, but instead retained a widespread nucleoplasmic distribution (Fig. 5B). We note this differential localization response to hinokiflavone treatment for the two sets of splicing proteins closely matches the observed inhibition of spliceosome formation at the A complex observed with *in vitro* splicing extracts (Figure 3B). In contrast with these changes in nuclear organization, we observed little or no effect of hinokiflavone treatment on either cytoplasmic structures, or on the localization of multiple cytoplasmic proteins (data not shown).

**Figure 5.**
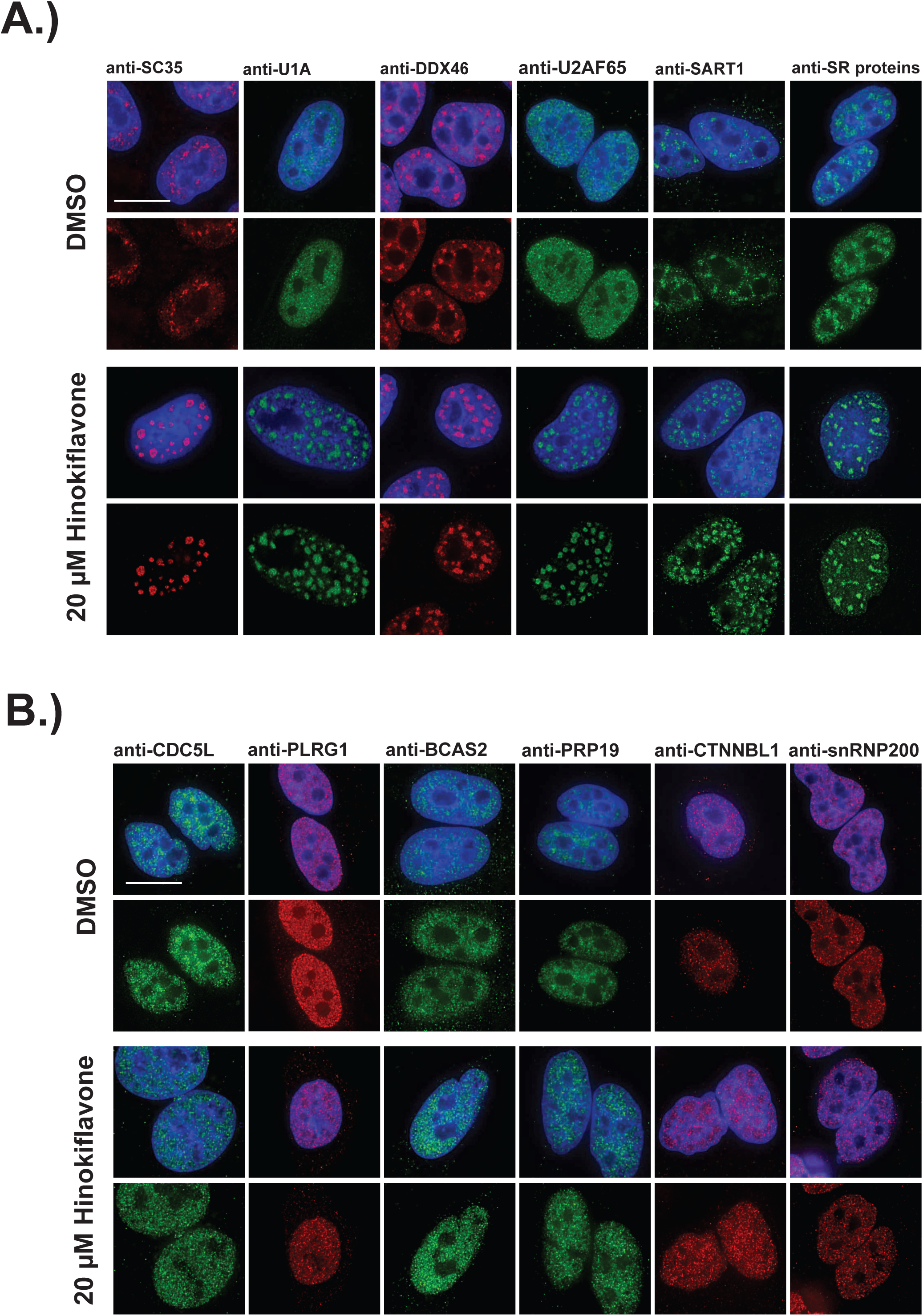
Changes in splicing speckles after treatment with hinokiflavone. HeLa cells incubated for 24h with either DMSO (control), or 20 μM hinokiflavone, were fixed and stained with antibodies for the following splicing factors: SC35, U1A, U2AF65, SART1, SR proteins, CDC5L, PLRG1, BCAS2, PRP19 and CTNNBL1. (A) Splicing factors involved in the early steps of spliceosome assembly are located in enlarged splicing speckles (B) Splicing factors that assemble after A complex formation are not accumulated in enlarged speckles and show a more diffuse nucleoplasmic distribution. Scale bars, 15 μm.

HeLa cells treated with 20 μΜ hinokiflavone for 24h showed a loss of Cajal bodies (CBs), which in control cells typically accumulate splicing snRNPs and nucleolar snoRNPs, but not other protein splicing factors. Seven CB components were investigated, i.e., coilin, SMN, TMG-cap, Y12, SNRPA1, CDK and Fibrillarin. Consistent with previous observations, in the control, DMSO-treated HeLa cells, coilin was enriched specifically in several CB foci, while SMN was observed in CBs in the nucleus and in punctate cytoplasmic structures. Components of splicing snRNPs, including TMG-cap, Y12 and SNRPA1, were located in both nuclear splicing speckles and CBs, while CDK and Fibrillarin localized in CBs and in the nucleolus (Figure 6). After treatment with hinokiflavone, bright CB foci were no longer observed, with coilin, SMN, TMG-cap, Y12 and SNRPA1 all relocalized and enriched in mega speckles. However, the cytoplasmic pool of SMN remained and appeared unaffected by hinokiflavone. Neither CDK, nor Fibrillarin, relocalized to the mega speckles, but the nucleoplasmic component of the staining pattern for both proteins was lost after treatment and only nucleolar CDK and Fibrillarin could be detected.

**Figure 6.**
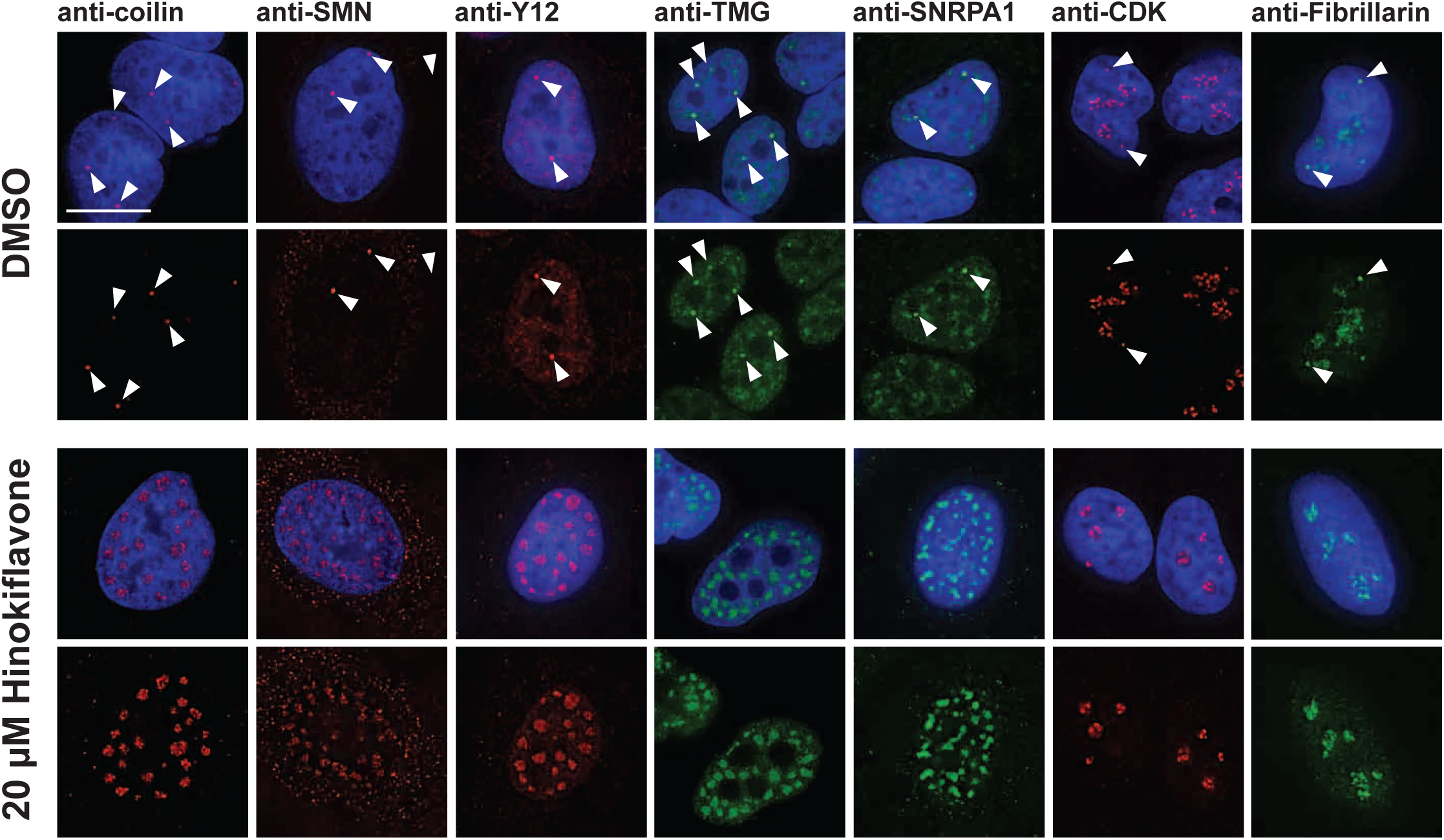
Hinokiflavone treatment leads to relocation of CB components to splicing speckles. HeLa cells were treated for 24h with either DMSO (control), or 20μM hinokiflavone and the fixed cells were stained with anti-coilin, anti-SMN, anti-Y12, anti-TMG, anti-SNRPA1, anti-CDK and anti-fibrillarin antibodies, respectively. Coilin, SMN, Y12 and TMG show relocation to enlarged splicing speckles in hinokiflavone treated cells. Arrowheads denote intact CBs. CDK and fibrillarin are only detected in nucleoli after treatment with hinokiflavone. Scale bars, 15 μm.

Disruption of CBs and relocalization of snRNP proteins is known to result from inhibition of transcription. We therefore compared the effect of hinokiflavone treatment with drugs that inhibit transcription, such as DRB, which inhibits transcription elongation by RNA polymerase II, using an assay to detect transcription sites by fluorescence microscopy. In contrast with the major inhibition of pre-mRNA synthesis caused by DRB (Figure 7A), we observe little or no change in RNA synthesis levels in HeLa cells, either 4h, or 8h, after treatment with either 10 μΜ, 20 μΜ, or 30 μΜ hinokiflavone (Figure 8A). However, we observed a dose dependent nuclear relocalization of snRPA1, coilin, TMG-cap and SMN into mega speckles and disruption of Cajal bodies (Figure 8B). While TMG-cap and snRPA1 relocated to mega speckles after treatment with DRB, coilin and SMN did not (Figure 7B). Coilin relocalized to the periphery of nucleoli and SMN dispersed as dots throughout the nucleoplasm. We conclude that the changes in subnuclear organisation induced by hinokiflavone are not likely to be indirect effects of inhibiting transcription.

**Figure 7.**
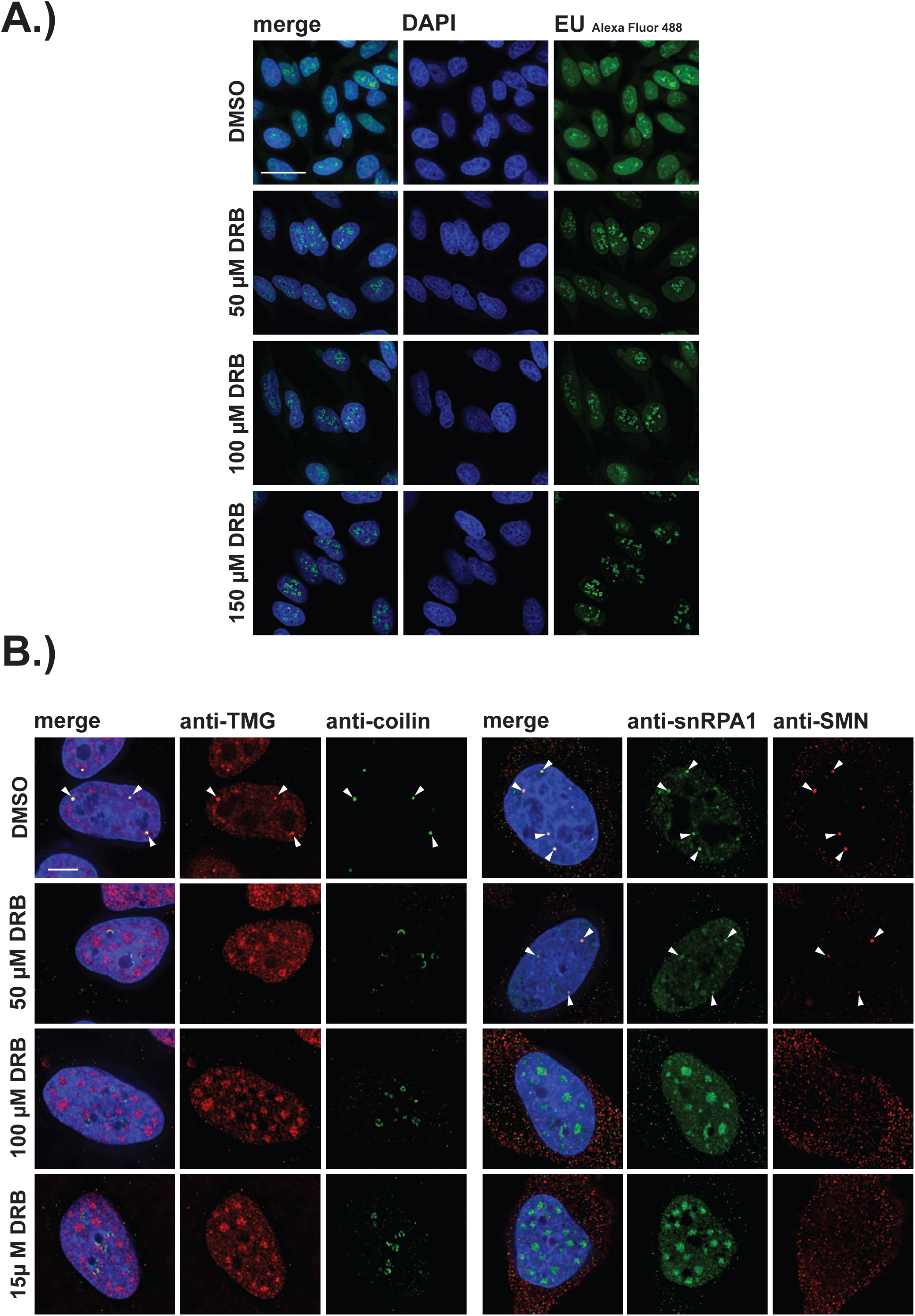
Inhibition of RNA polymerase II transcription and disruption of Cajal bodies by DRB. HeLa cells were incubated for 4h with DRB before cells were incubated with EU that was incorporated into newly synthesized RNA for 20 min. (A) Cells were fixed and labeled RNA was detected by fluorescence microscopy. Scale bar, 40 μm. (B) HeLa cells were incubated with either DMSO (control), or with 50, 100 or 150 μΜ DRB for 4h and stained with anti-coilin, anti-TMG, anti-SMN and anti-snRPA1 antibodies, respectively. Arrowheads denote intact CBs. Scale bar, 6.5 μM

**Figure 8.**
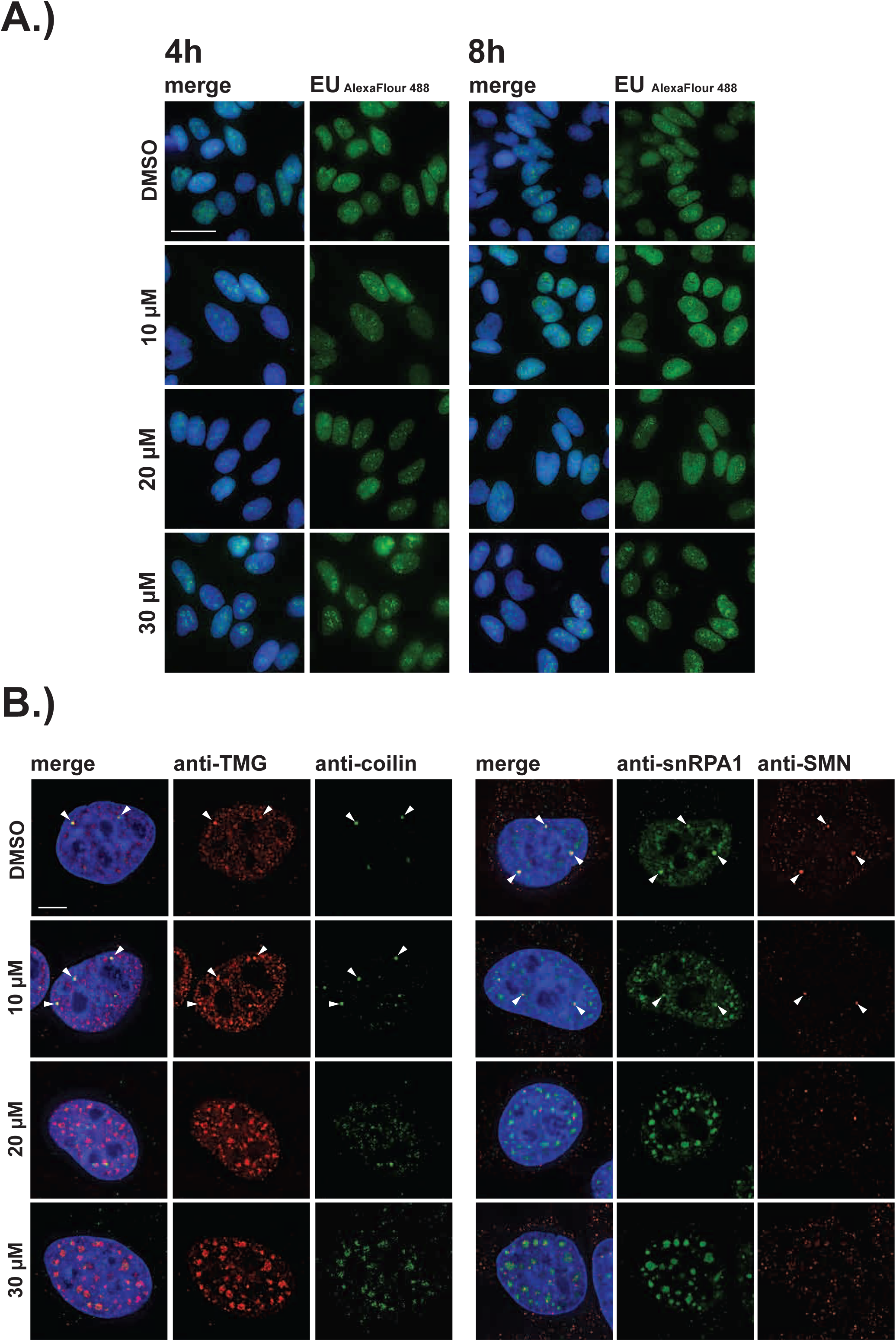
Disruption of Cajal Bodies, but no inhibition of polymerase II transcription after 8h of treatment with hinokiflavone. (A) HeLa cells were treated with either DMSO (control), or with 10, 20 or 30 μM hinokiflavone for either 4h, or 8h, then cells were incubated with EU that was incorporated into newly synthesized RNA for 20 min. Cells were fixed and labeled RNA was detected by fluorescence microscopy. Scale bar, 40 μm. (B) HeLa cells were incubated with either DMSO (control), or with 10, 20 or 30 μM hinokiflavone for 8h then fixed and stained with anti-coilin, anti-TMG, anti-SMN and anti-snRPA1 antibodies, respectively. Arrowheads denote intact CBs.

### Hinokiflavone promotes nuclear relocalization of SUMO

Due to the unexpected changes in the localization of coilin and SMN induced by hinokiflavone, we next tested whether the localization of other nuclear proteins, including PML, the cleavage stimulation factor subunit 2 (CSTF2), SUMO1 and SUMO2/3, was affected. We observe loss of PML-containing nuclear bodies (PML-NBs), after treatment with hinokiflavone, with PML relocalized into a pattern of dots, located at the periphery of the enlarged speckles containing splicing factors (Figure 9A). The diffuse nucleoplasmic localization of CSTF2 seen in control cells was lost, with CSTF2 also relocated to dots at the periphery of the mega speckles (Figure 9B).

**Figure 9.**
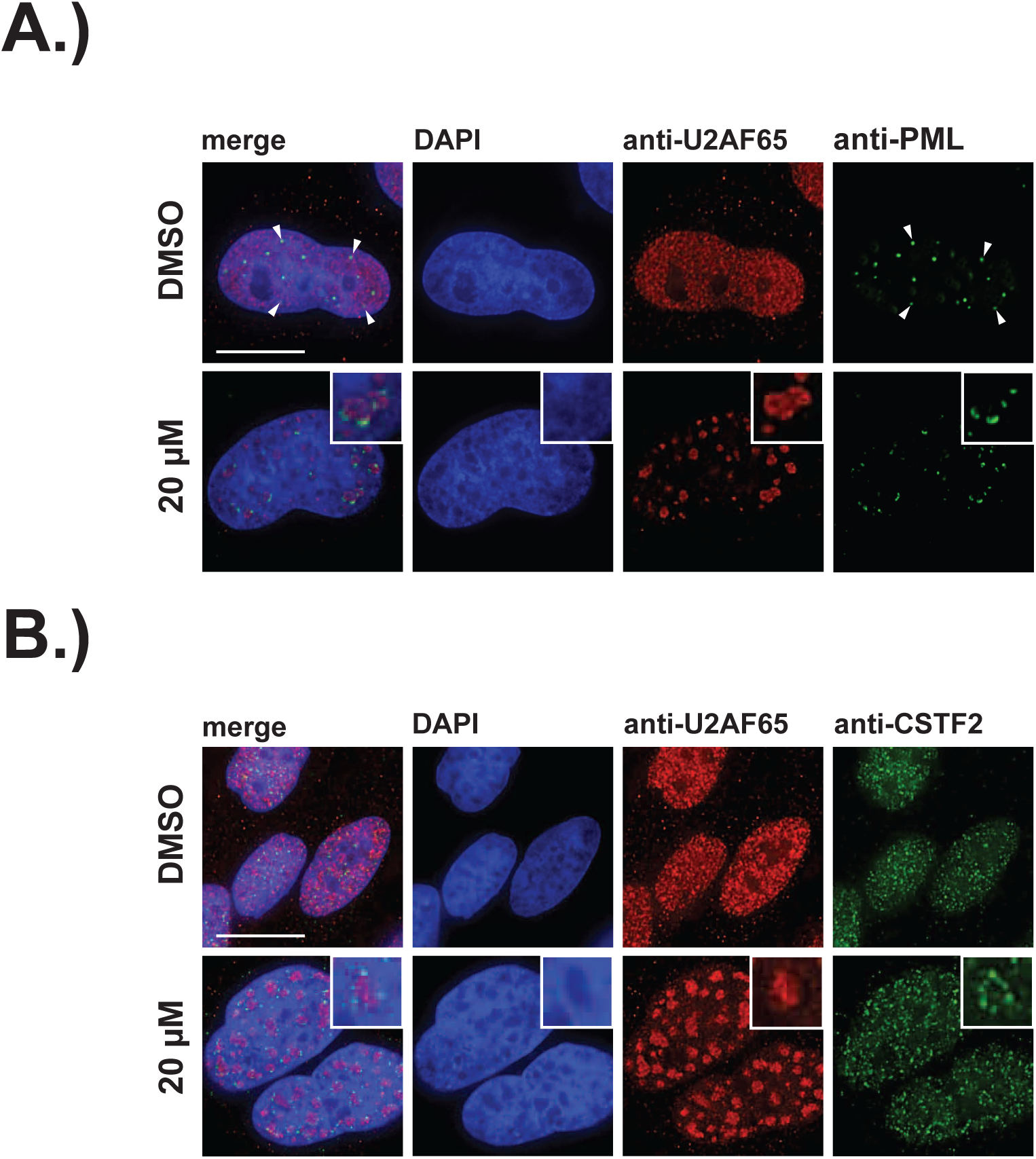
Hinokiflavone treatment leads to the relocation of nuclear proteins to the periphery of enlarged splicing speckles. (A) HeLa cells were treated for 24h with either DMSO (control), or 20 μM hinokiflavone and the fixed cells were stained with either anti-CSTF2, or anti-PML antibodies. Arrowheads denote PML bodies (B). Costaining with anti-U2AF65 antibodies showed that both CSTF2 and PML relocate to the periphery of enlarged splicing speckles. Scale bars, 15 μm.

In the DMSO treated control cells, SUMO2/3 showed diffuse nucleoplasmic staining and bright foci, while SUMO1 showed both nuclear membrane staining and concentration in nucleoplasmic foci. Surprisingly, we observed both SUMO1 and SUMO2/3 staining also relocalized to mega speckles in hinokiflavone-treated cells (Figure 10A). This was also seen in HeLa cells treated for 2h with 10 μΜ, 20 μΜ and 30 μΜ hinokiflavone, then stained with antibodies specific for either SUMO1, SUMO2/3 or SFRS2 (SC35). Already after this short, 2h exposure to hinokiflavone, all three markers show co-localization in speckles (Figure 10B).

**Figure 10.**
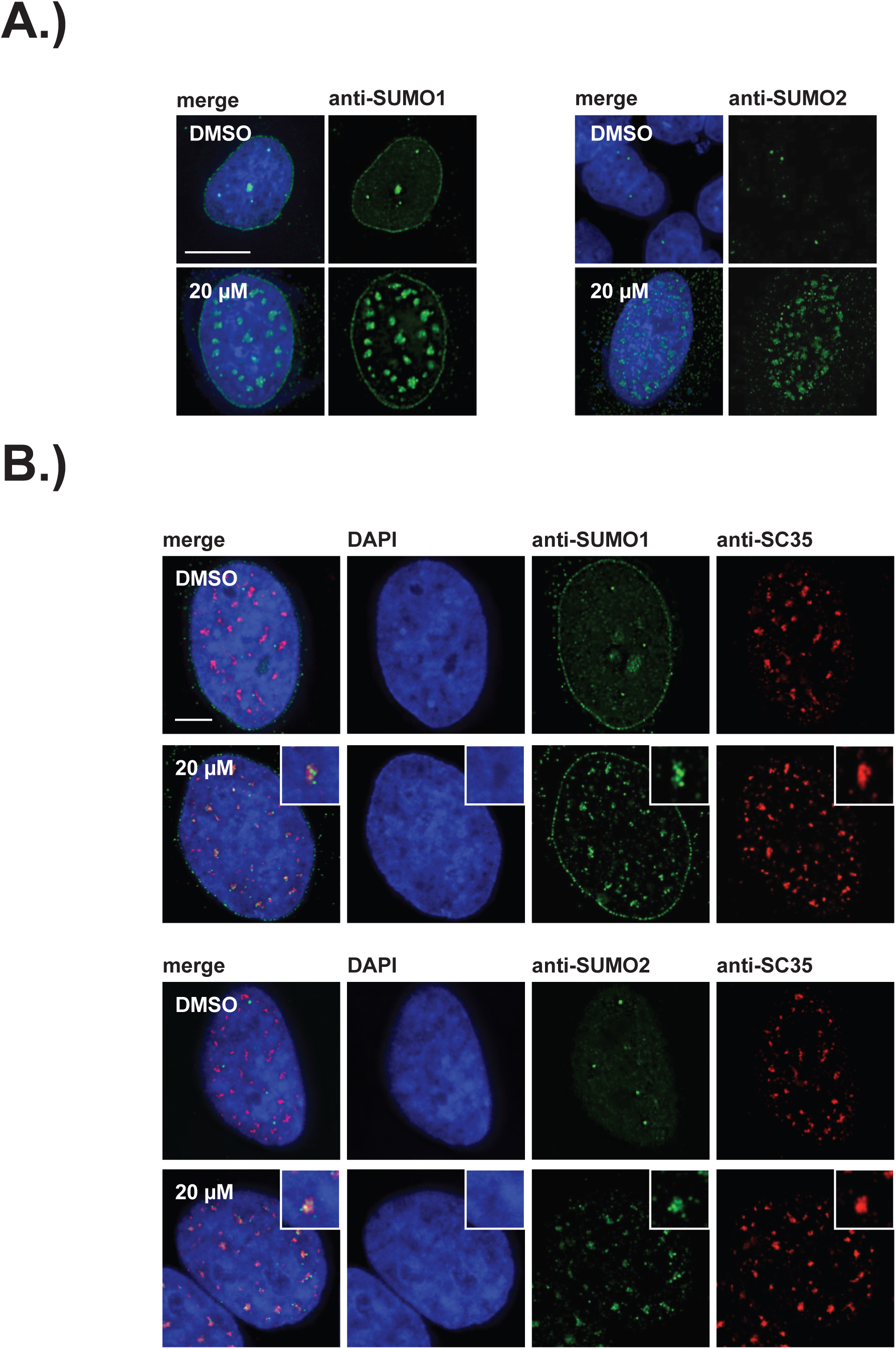
SUMO1 and SUMO2/3 relocalize to enlarged splicing speckles in the presence of hinokiflavone. (A) HeLa cells were treated for 24h with either DMSO, or 20 μM hinokiflavone and the fixed cells were stained with either anti-SUMO1, or anti-SUMO2/3 antibodies. Both SUMO1 and SUMO2/3 accumulated in the enlarged splicing speckles formed after treatment with hinokiflavone. (B) Treatment of HeLa cells with 20 μΜ hinokiflavone for 2h. Co-staining with either anti-SUMO1, or anti-SUMO2/3 and anti-SC35 antibodies showed that both SUMO1 and SUMO2/3 accumulate in enlarged splicing speckles. Scale bars, 15 μm.

### Hinokiflavone increases levels of SUMOylated proteins

Following the observed relocalization of SUMO1 and SUMO2/3 to mega speckles, we next tested whether hinokiflavone treatment altered the protein SUMOylation pattern in cells. For this, HEK293 cells were treated for 24h with either DMSO (control), or with 10 μM, 20 μM, or 30 μM hinokiflavone, then the cells were harvested and total cell lysate separated by SDS-PAGE and analyzed by immunoblotting (Figure 11A). This shows a clear increase in the accumulation of high molecular weight, SUMO1 and SUMO2/3- modified proteins in the hinokiflavone treated extracts, as compared with the DMSO control. Interestingly, parallel immunoblotting analysis for the related peptide modifiers ubiquitin and NEDD8, showed no increase in their levels of protein conjugation after hinokiflavone treatment, but rather a modest decrease (Figure 11B). This demonstrates a remarkably selective effect of hinokiflavone for promoting specifically increased levels of protein SUMOylation *in cellulo*. We also observed an increase in protein SUMO2/3 SUMOylation after hinokiflavone treatment of HeLa and NB4 cells (Supplementary Figure 4).

**Figure 11.**
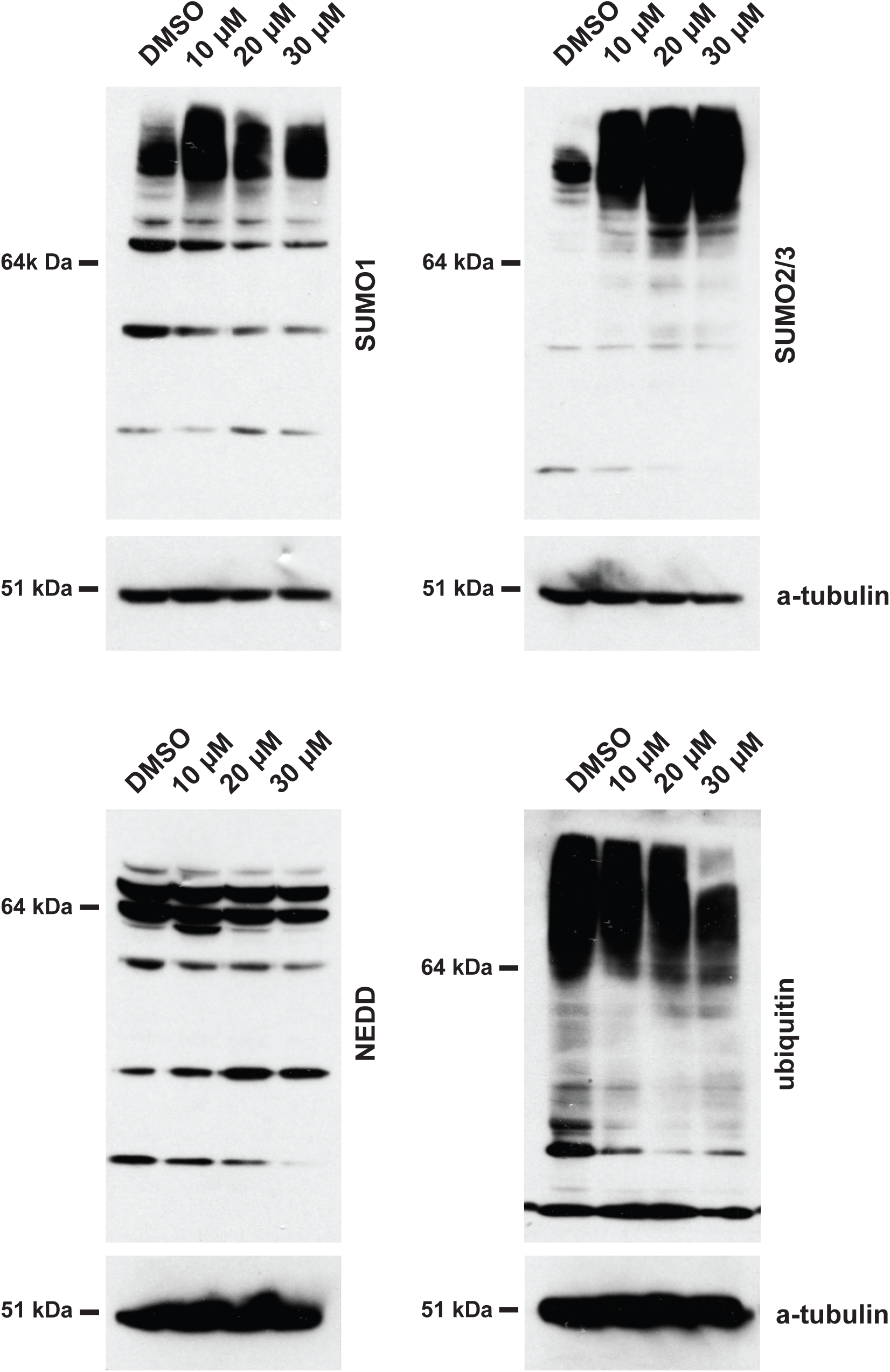
Treatment of HEK293 cells with hinokiflavone leads to an increase in SUMOylated proteins. HEK293 cells were treated with either DMSO, or with 10 μM, 20 μM, or 30 μM hinokiflavone for 24h before cells were lysed in 1x LDS buffer. Samples were separated on SDS-PAGE and transferred to membranes. After probing with antibodies specific for SUMO1, SUMO2/3, ubiquitin, and NEDD8, labeled proteins were visualized using chemiluminescence, showing a specific accumulation of poly-SUMOylated proteins after hinokiflavone treatment. The membranes were also probed to detect alpha-tubulin as a negative control (bottom panels).

We next analyzed the effect of hinokiflavone on protein SUMOylation levels in the nuclear extracts used for *in vitro* splicing reactions. For this, splicing reactions were carried out using HeLa cell nuclear extract and treated with either DMSO (control), or with hinokiflavone, at a final concentration of either 100 μM, 300 μM, or 500 μΜ. After incubation at 30^°^C for 90 min, the splicing reactions were separated by SDS-PAGE and immuno-blotted to detect either SUMO1, SUMO2/3 or SFRS1, the latter acting as a loading control (Figure 12). In the presence of hinokiflavone we observed a major increase in the levels of high molecular weight, SUMO1 and SUMO2/3 modified proteins, when compared with the DMSO control (Figure 12). Furthermore, this increase in protein SUMOylation levels is hinokiflavone concentration-dependent.

**Figure 12.**
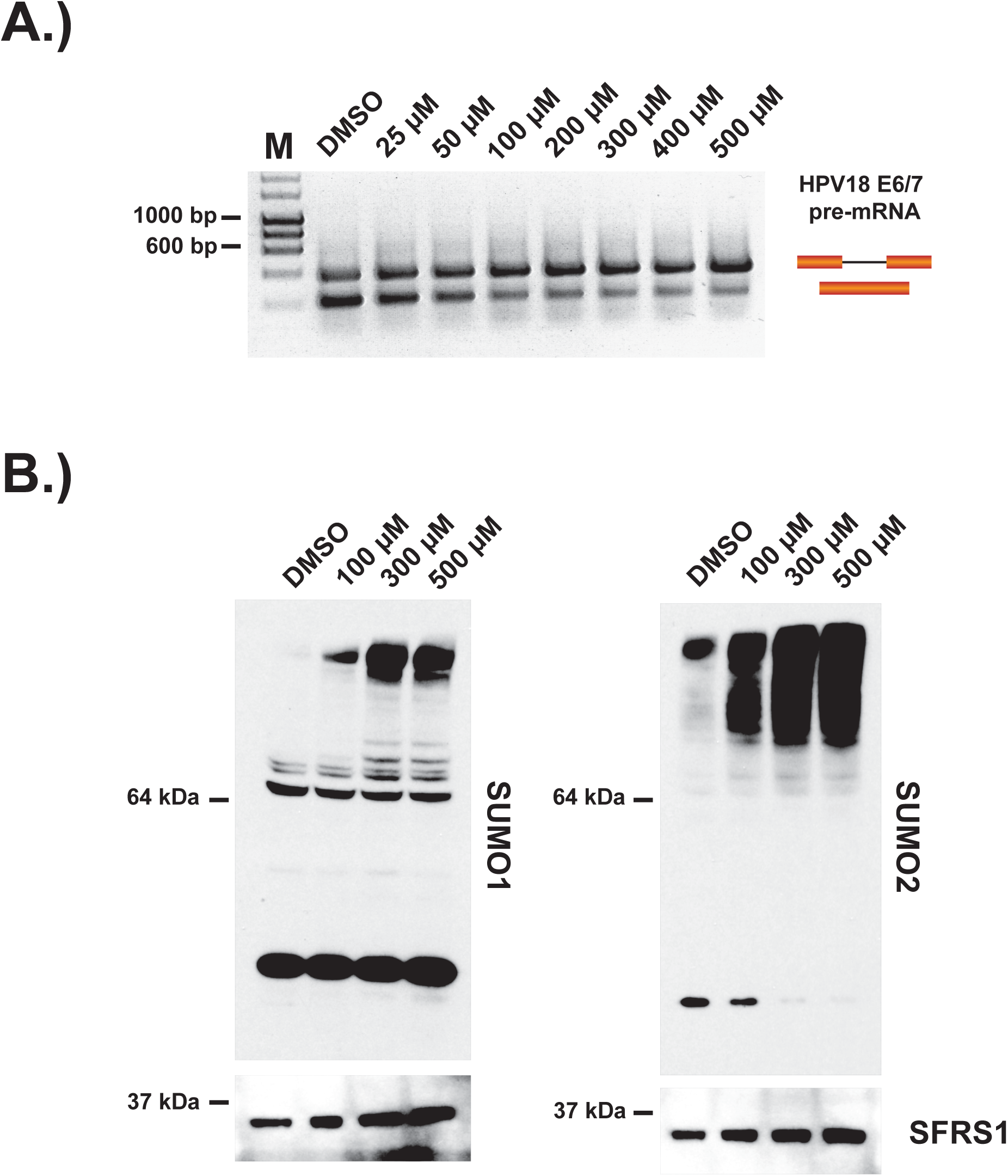
Incubation of *in vitro* nuclear splicing extracts with hinokiflavone leads to an increase in SUMOylated proteins. HeLa nuclear extracts reactions were incubated *in vitro* under splicing conditions with either DMSO (control), or with increasing concentrations from 25-500 μM hinokiflavone. (A) Evaluation of the lowest concentration at which hinokiflavone inhibits pre-mRNA splicing *in vitro*. (B) Proteins were extracted and size separated by SDS-Page, transferred to membranes and probed using either anti-SUMO1, anti-SUMO2/3, or anti-SRSF1 antibodies and visualized using chemiluminescence, showing a specific accumulation of hyper-SUMOylated proteins after hinokiflavone treatment. The membranes were also probed to detect SFRS1 as a loading control (bottom panels).

### Hinokiflavone inhibits SENP1 activity

Protein SUMOylation is a reversible modification. The level of SUMOylated protein thus reflects the balance between the rate of SUMO conjugation by SUMO E3 ligases and the rate of SUMO deconjugation, catalyzed by sentrin-specific proteases (SENPs). We therefore hypothesized that the observed increase in protein SUMOylation caused by hinokiflavone, both *in cellulo* and *in vitro*, could result from an inhibition of SENP activity. To test this hypothesis, we performed *in vitro* SENP assays, using a purified, catalytically active fragment of the SENP1 protein (aa 415-643), which was expressed in *E.coli* (Figure 13A). We compared SENP1 activity when incubated in the presence of either DMSO (control), or with either 500 μM hinokiflavone, or the four other biflavones previously tested for their ability to inhibit splicing (i.e., amentoflavone, cupressuflavone, isoginkgetin and sciadopitysin). The SENP assays were carried out as described in experimental procedures and then the reactions separated using a 4-12% Bis-Tris PAGE gel and proteins were visualized by staining with coomassie blue (Figure 13A). This showed a clear inhibition of SENP1 activity by hinokiflavone (lane 5), as compared with the DMSO control (lane 4). Interestingly, SENP1 inhibition was also seen with some of the other biflavones, with the degree of SENP inhibition correlating with their ability to inhibit pre-mRNA splicing *in vitro* (cf. Figure 1B). Thus, hinokiflavone and amentoflavone show the greatest inhibitory effect *in vitro* on both splicing and SENP1 activity.

**Figure 13.**
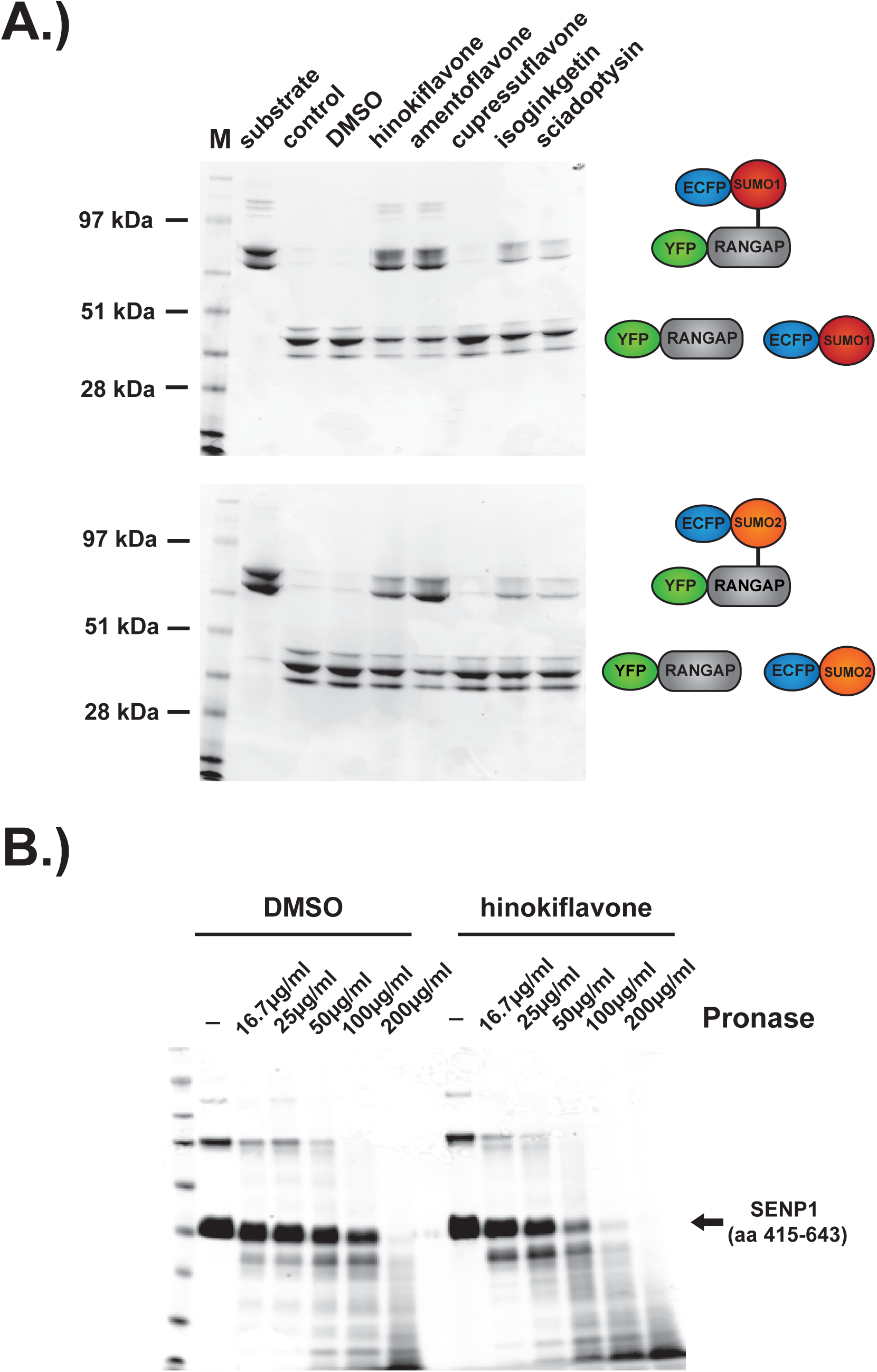
Biflavones inhibit SENP 1 *in vitro*. The effect of 500 μM hinokiflavone, amentoflavone, cupressuflavone, isoginkgetin and sciadopitysin on the isopeptidase activity of a highly purified fragment of catalytically active SENP1 (comprising aa 415-643), that had been expressed in *E. coli*, was determined by *in vitro* gel-based activity assay (A). Incubation of the catalytically active SENP1 fragment with either DMSO or 500 μM hinokiflavone, before the samples were digested with different concentrations (from ∼16 – 200 micrograms/ml) of the proteinase pronase. SENP1 showed an increased sensitivity to protease digestion in the presence of hinokiflavone.

The drug affinity responsive target stability (DARTS) assay provides a convenient way of testing whether a compound binds to putative target proteins (^17,18^). We therefore used the DARTS approach to evaluate whether hinokiflavone can bind to recombinant SENP1. For this assay, the purified, catalytically active SENP1 protein fragment was incubated with either DMSO (control), or 500 μM hinokiflavone, before the samples were subjected to limited digestion with varying concentrations of pronase, from ∼16 – 200 micrograms/ml (Figure 13B). This showed that in the presence of hinokiflavone, the sensitivity of the SENP1 fragment to protease digestion increased, indicating that hinokiflavone directly interacts with SENP1.

In summary, these data show that a structurally related group of biflavone compounds can inhibit purified SENP1 SUMO protease *in vitro* and this correlates with their potency as *in vitro* inhibitors of pre-mRNA splicing. The strongest inhibitor, hinokiflavone, was further shown to interact with SENP1 using the DARTS assay. These data support the hypothesis that cells treated with hinokiflavone are prevented from de-conjugating SUMO.

### Identification of protein SUMOylation targets affected by hinokiflavone

Having identified hinokiflavone as a SUMO protease inhibitor *in vitro*, which causes the accumulation of poly-SUMOylated proteins *in cellulo*, we next sought to identify protein targets whose level of SUMO modification increases in cells treated with hinokiflavone. For this analysis we used a recently described, quantitative, SILAC proteomics approach in HEK293 SUMO2^T90K^ cells^19^. Thus, HEK293 SUMO2^T90K^ cells, which express the his-tagged SUMO2^T90K^ mutant, were treated for 8h with either DMSO (control), or with 20 μM hinokiflavone and lysates prepared. SUMO2^T90K^-conjugated target proteins in the lysates were affinity purified under denaturating conditions using the His-tag. After cleavage with endonuclease LysC, an antibody specific for the diGly-Lys peptide, which is diagnostic of SUMO modification, was used to enrich for peptides including lysine residues that had been SUMO2^T90K^-modified.

This analysis identified 961 SUMO2-modified lysine residues in 553 different proteins (Supplementary Table 1). Of these, twenty-four lysine residues showed more than a five fold increase in the level of SUMO2 modification in the hinokiflavone-treated cells, when compared with the DMSO control. Interestingly, eleven of these sites are located in six proteins that are components of the U2 snRNP, i.e., PRPF40A, SF3B2, SF3A2, SNRPD2, U2SURP and SF3B1 (Figure 14A). These six proteins have previously been shown to interact, as illustrated in Figure 14B. The most dramatic effect of hinokiflavone was seen for the U2 snRNP associated protein PRPF40A, which increased SUMO2 modification levels at four lysine residues, i.e. K241, K375, K517 and K707, including a remarkable increase in the level of SUMO2 at K241 of ∼145 fold in the hinokiflavone-treated cells.

**Figure 14.**
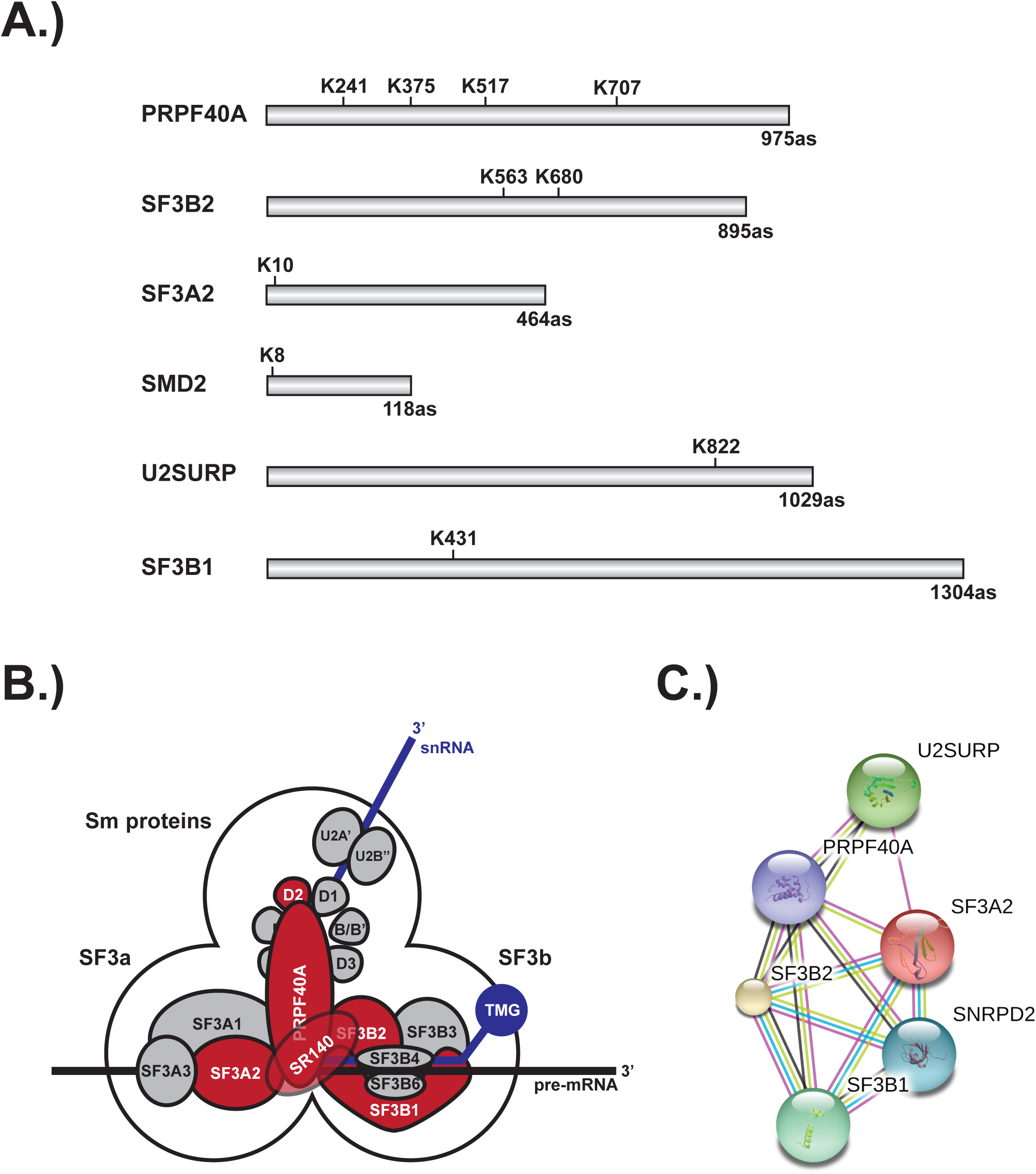
Schematic representation of splicing factors identified as SUMO2 target proteins. (A) Lysine residues in PRPF40A, SF3B2, SF3A2, SMD2, SR140 and SF3B1 that show an upregulated SUMO2 SUMOylation in HEK293 cells treated for 8h with 20 μΜ hinokiflavone are shown. (B) Schematic representation of the SUMO2 modified U2 snRNP components, which are coloured in red. (C) STRING network diagram showing interactions between the U2 snRNP proteins detected as having SUMO2 modification levels upregulated > 5 fold in cells after hinokiflavone treatment.

To validate these MS data, we performed immuno-blotting experiments, using an antibody specific for PRPF40A, with extracts prepared from HEK293 cells treated for 24h with either DMSO (control), or with 10 μM, 20 μM, or 30 μM hinokiflavone (Figure 15A). This showed that the level of SUMOylated PRPF40A increased following incubation of cells with hinokiflavone. In addition, immunofluorescence analysis using the same antibody showed that after hinokiflavone treatment PRPF40A accumulated in the mega speckles (Figure 15B).

**Figure 15.**
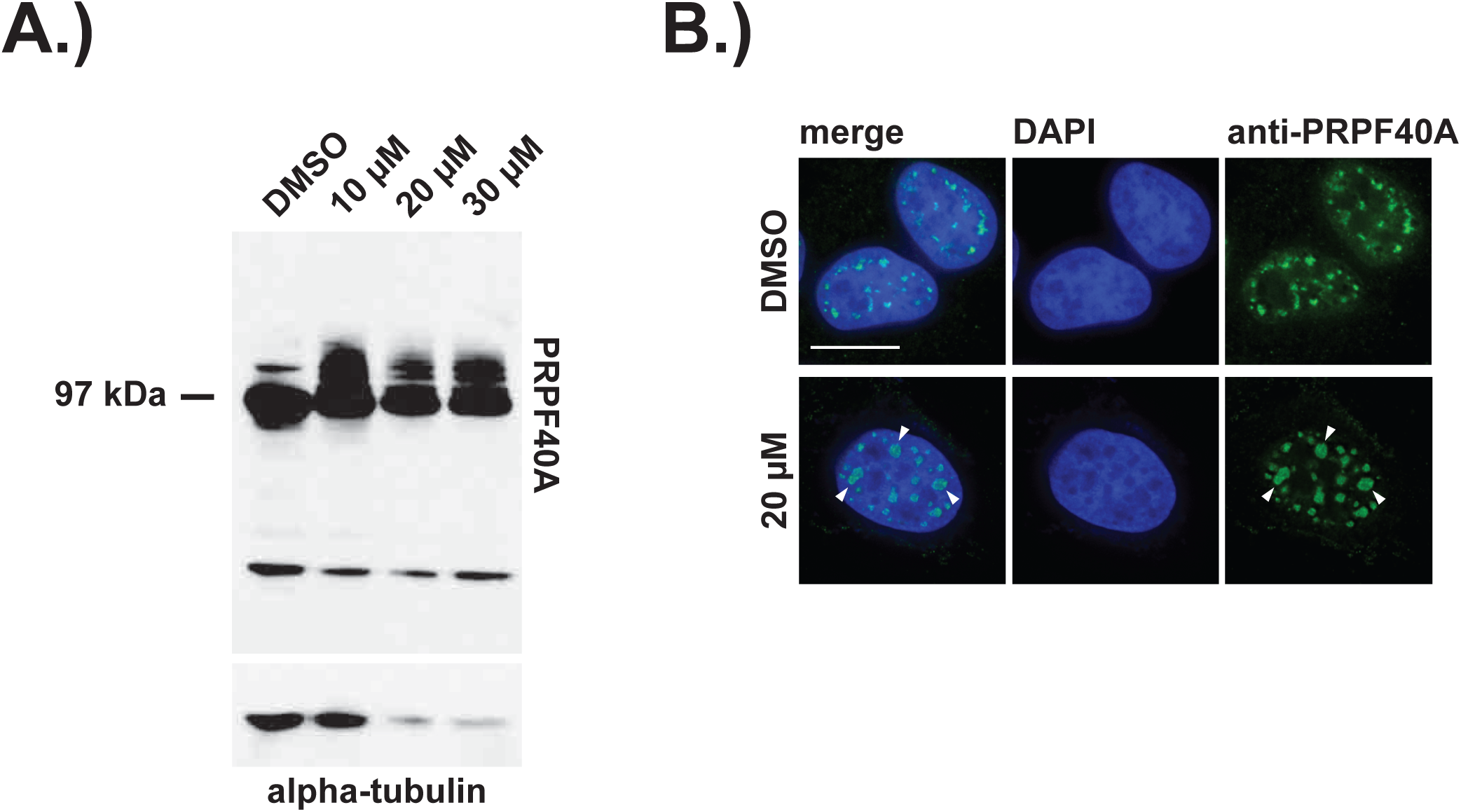
Confirmation of PRPF40A as a SUMO target protein. (A) HEK293 cells were treated with either DMSO (control), or with 10 μΜ, 20 μΜ or 30 μΜ of hinokiflavone for 24h, then total cell lysates were size-separated by SDS-PAGE, transferred to membranes and probed using the anti-PRPF40A antibody and visualized using chemiluminescence. Arrows indicate SUMO2 modified proteins. (B) Immunofluorescence analysis shows that hinokiflavone treatment leads to the relocation of PRPF40A to mega speckles in HeLa cells.

### SUMOylated splicing factors are found in the insoluble protein fraction after treatment with hinokiflavone

HEK293 cells stably expressing GFP-PRPF40A were established and the behaviour of GFP-PRPF40A in the presence of hinokiflavone was tested. Like endogenous PRPF40A, GFP-PRPF40A relocated to mega-speckles after hinokiflavone treatment (Figure 16A). This cell line was then used to examine if the SUMO2 SUMOylation of PRPF40A changed the interactions between PRPF40A and other splicing factors. GFP-PRPF40A expressing cells were treated with either 20 μM hinokiflavone, or DMSO (control), for 8h before cells were lysed with Co-IP buffer. GFP-tagged proteins were immunopreciptated using GFP-Trap beads and analyzed by western blotting (Figure 16B). Immunopreciptation of PRPF40A co-isolates SF3B2, but, as expected, not PRP19. However, SUMOylated PRPF40A and SF3B2 could only be detected in the non-soluble fraction of hinokiflavone treated cells. These data are consistent with hinokiflavone promoting hyper-SUMOylation of splicing factors, resulting in their forming insoluble aggregates and/or other forms of high molecular weight complexes.

**Figure 16.**
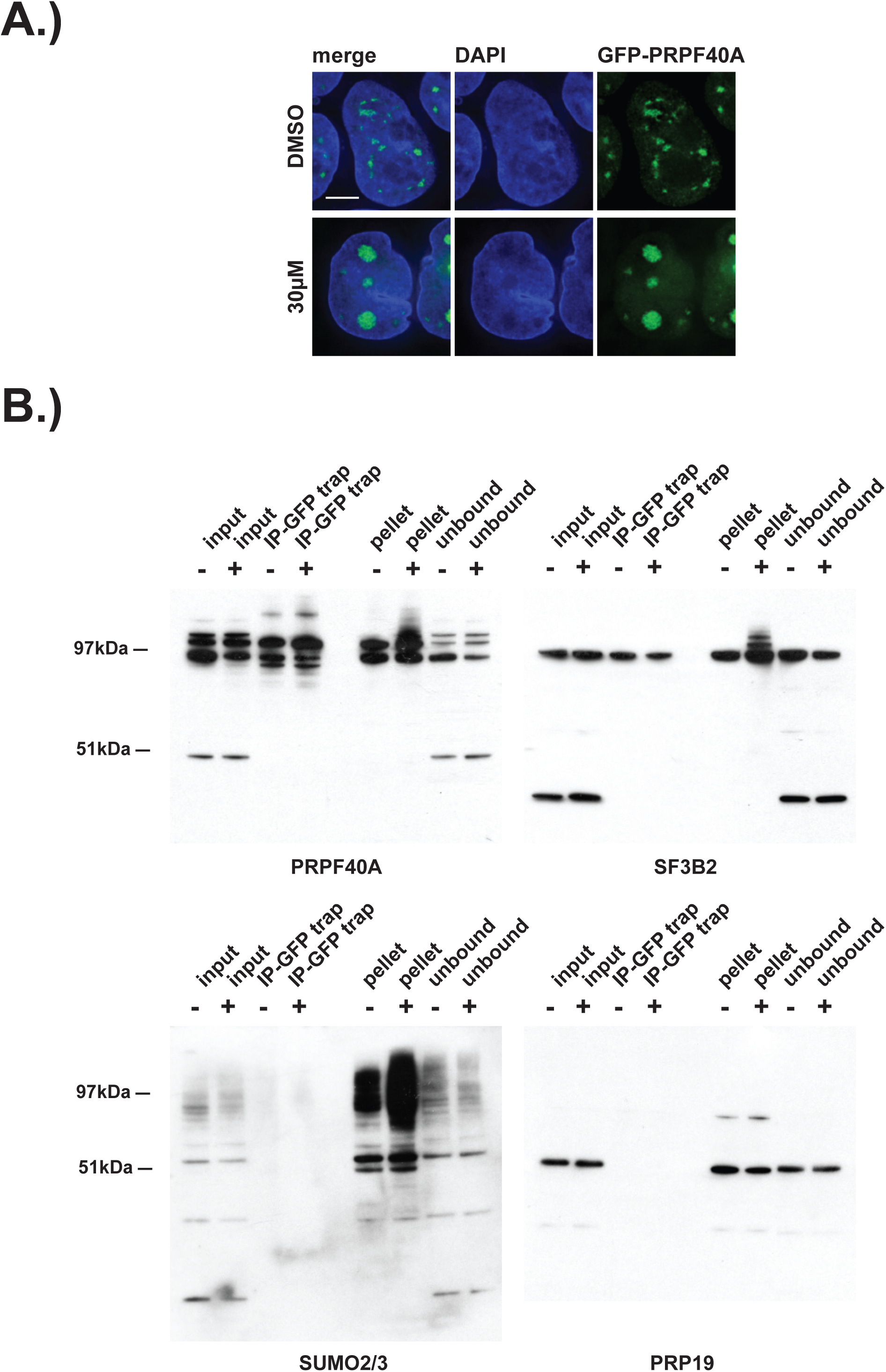
SUMOylated PRPF40A and SF3B2 are increased in the insoluble fraction of HEK293 cell lysates. HEK293 cells were treated with either DMSO (−) as a negative control, or with 20 μM hinokiflavone (+), for 8h, then cells were lysed and analyzed by co-immunoprecipitation (Co-IP). The input, IPs, pellets and unbound fractions of both the control and hinokiflavone treated cells were size separated by SDS-Page, transferred to membranes and probed using anti-PRPF40A, anti-SF3B2, anti-SUMO2/3 and anti-PRP19 antibodies and visualized using chemiluminescence.

## Discussion

In this study we have identified the plant biflavone, hinokiflavone, to be a novel modulator of pre-mRNA splicing activity, both *in vitro* and *in cellulo*. Hinokiflavone blocks splicing of pre-mRNA substrates *in vitro* by inhibiting spliceosome assembly after formation of the A complex. Multiple human cell lines treated with hinokiflavone show changes in alternative splicing patterns and altered nuclear organization of splicing factors required for A complex formation, resulting in enlarged nuclear splicing speckles enriched in SUMO1 and SUMO2 and concomitant disruption of Cajal bodies (CBs). Hinokiflavone treated cells show dose and time-dependent cell cycle arrest phenotypes, along with varying degrees of apoptosis, the latter effect seen most strikingly in the promyelocytic NB4 cell line that expresses the PML-RARalpha fusion protein.

Hinokiflavone treated cells accumulate hyper-SUMOylated proteins, which we hypothesise results from hinokiflavone inhibiting de-SUMOylation. Consistent with this, we show that *in vitro* hinokiflavone can inhibit the catalytic activity of a purified, *E. coli* expressed fragment of the SUMO protease SENP1. Using a quantitative, mass spectrometry-based assay we identified protein targets and mapped lysine residues showing increased levels of SUMO2 modification in hinokiflavone treated cells. The major target proteins were enriched in pre-mRNA splicing factors, in particular, six components of the U2 snRNP spliceosome subunit that is required for A complex formation. This included four lysine residues in the U2 snRNP protein PRPF40A, with K241 showing a remarkable increase of >140 fold in levels of SUMO2 modification after hinokiflavone treatment. Our data thus provide multiple lines of evidence linking SUMO modification of proteins in the pre-mRNA splicing machinery with downstream effects on alternative splicing, nuclear organization and cell cycle progression.

To date, the best studied group of splicing modulators are the SF3B1 inhibitors, such as Spliceostatin A, which either have been isolated from the broth of bacteria, or else are synthetic derivatives of these compounds (^12-14^). Several other natural compounds isolated from plants have also been shown to affect pre-mRNA splicing, including the flavones apigenin^20^ and luteolin^21^ and the biflavone isoginkgetin^16^, but in comparison with the SF3B1 inhibitors, very little is known about either their targets, or their mode of action. In this study we compared the ability of the biflavones amentoflavone, cupressuflavone, hinokiflavone and sciadopitysin, to modify splicing, either *in vitro* or *in cellulo*, comparing them with the previously described biflavone splicing inhibitor isoginkgetin. Our analysis showed that within this group of compounds hinokiflavone was the most potent biflavone modulator of splicing.

Hinokiflavone is a biapigenin found in many different plant families^22^. Previous studies, using *in silico* screens, have suggested several different proteins as potential targets for hinokiflavone, including the prostaglandin D2 synthetase^23^ and the matrix metalloproteinase-9^24^. We also tested the ability of hinokiflavone to inhibit a large panel of purified kinases *in vitro* and observed that it had either no, or only weak, non-specific inhibitory effects on any of the 120 different kinases tested (data not shown). In contrast, however, using direct biochemical assays, we identify here that hinokiflavone is a SUMO protease inhibitor. Specifically, we show that, (a) hinokiflavone inhibits the catalytic activity of a highly purified, *E. coli* expressed fragment of the SUMO protease SENP1 and (b) that treatment of multiple human cell lines with hinokiflavone resulted in a dramatic increase in the levels of SUMO1 and SUMO2/3 modified proteins. The effect in hinokiflavone treated cells appears to be specific to SUMO modification because we detected little or no parallel increase in the levels of proteins linked with related post-translational peptide modifiers, i.e. either ubiquitin, or NEDD8. Accumulation of high molecular weight SUMOylated proteins was also observed *in vitro* when HeLa nuclear extracts were incubated with hinokiflavone.

We characterized in more detail the stimulation of SUMO modification in cells treated with hinokiflavone, using a recently developed, quantitative, MS-based proteomics assay in HEK293 SUMO2^T90K^ cells^19^. We identified 961 SUMO2 modified lysine residues in 553 target proteins and showed that 24 of these lysine residues increased SUMO2 levels >5 fold after hinokiflavone treatment. Interestingly, this unbiased assay independently linked the effect of hinokoflavone with the pre-mRNA splicing machinery, with 11 of the lysine residues showing the highest increase in SUMO2 modification located specifically in 6 U2 snRNP proteins.

We note that U2 snRNP is required for assembly of the pre-splicing A complex and that hinkoflavone prevents spliceosome assembly *in vitro* proceeding beyond formation of the A complex. It is possible, therefore, that one or more U2 snRNP proteins are transiently SUMOylated during the spliceosome assembly cycle and must be de-SUMOylated for assembly to proceed beyond the A complex. For example, we speculate this could be involved in a proof reading mechanism that ensures accurate selection of 5’ and 3’ splice sites before proceeding to form a catalytically active complex. This would provide a potential mechanism linking our observations that hinokiflavone both inhibits protein de-SUMOylation and prevents spliceosome assembly *in vitro* proceeding beyond the A complex. This would be consistent with our additional observations by fluorescence microscopy that only splicing factors involved in the early steps of the splicing process are enriched in the mega-speckles that are formed in hinokiflavone treated cells, while other splicing factors associated with later steps in spliceosome assembly remain mostly diffusely distributed throughout the nucleus.

Our current data mapping lysine residues in U2 snRNP proteins that are SUMO modified and which show enhanced SUMO2 modification after hinokiflavone treatment, are consistent with previous data suggesting a potential link between splicing and protein SUMOylation. For example, Pelisch et al., reported that the ser/arg-rich non-snRNP protein splicing factor SRSF1 regulates protein SUMOylation and interacts with the SUMO E3 ligase PIAS1^25^. Interestingly, SRSF1 was originally identified independently as both an essential splicing factor *in vitro* and also as a factor involved in the control of alternative splice site choice in cells^26,27^. Detailed biochemical studies have shown that SRSF1 is multifunctional, with important roles during early steps in spliceosome assembly, leading to formation of the A complex.

Two recent proteomics studies that systematically identified proteins that are SUMO2 modified in cells, using high throughput, mass spectrometry-based proteomics methods, reported splicing factors amongst the many targets detected in HEK293^19^ and HeLa cells^28^. Both these studies identified PRPF40A as a SUMO2 target protein. In addition Tammsalu et al., identified SNRPD2 and Hendriks et al., identified the U2 snRNP components SF3A2, SF3B1 and SF3B2 as targets for SUMO2 modification. Furthermore, in a recent study identifying the SENP inhibitor SI2, it was reported that blocking SENP activity increase the levels of SUMOylation of multiple proteins, including the splicing factors USP39, SF3B1 and PRPF40A^29^. While in this latter study the authors did not map the specific lysine residues that were modified by SUMO2 and none of these three studies investigate whether SUMO modification, or inhibition of SENP activity, affects pre-mRNA splicing, it is nonetheless interesting that here we also find increased SUMO2 modification of the same U2 snRNP proteins, SF3B1 and PRPF40A, upon inhibition of SENP activity by hinokiflavone. Indeed, our data show that PRPF40A had the highest overall increase in SUMO2 modification of all the SUMOylated proteins detected after treatment with hinokiflavone, including one lysine residue (K241) that increased ∼145 fold.

PRPF40A is a conserved spliceosome protein, present throughout eukaryotes from yeast to mammals. It was identified as a component of the U2 snRNP spliceosome subunit and suggested to have a role in mediating interactions between the separate 5’ and 3’ splice sites on pre-mRNAs^30^. It is intriguing that all the splicing factors that we detect here as showing the highest increase (>5 fold) in SUMO2 levels in cells treated with hinokiflavone, i.e., SF3A2, SF3B1, SF3B2, SNRPD1 and U2SURP, are either core components of, or associated with, U2 snRNPs and have been shown to interact with each other^30^. Several reversible, post-translational modifications, including phosphorylation, ubiquitination, methylation and acetylation, were previously reported to be important for the assembly and the disassembly of the spliceosome^31^. Our present data suggest that reversible SUMO modification may also be an important feature of mechanisms affecting spliceosome assembly and alternative splice site choice.

Previous studies using the SF3B1-targeted splicing inhibitors have shown that they are potent growth inhibitors of many different cancer cell lines (^32,33^). When used in combination with Bcl-2 inhibitors, these SFB31 inhibitors have been demonstrated to induce apoptosis in small lung cancer cells, chronic lymphocytic leukemia cells and head and neck cancer cells (^34-36^). This has been mainly attributed to their ability to change the alternative splicing of MCL1 pre-mRNAs. Indeed, maeyamcin and spliceostatin A showed the greatest effect on exon 2 skipping of MCL1 when compared with their effects on alternative splicing of 34 other genes important for cell proliferation and apoptosis^37^. Here we show that, in common with the previously demonstrated effect of these SF3B1 inhibitors, hinokiflavone treatment also changes the alternative splicing of MCL1 pre-mRNA to favour the production of the proapoptotic isoform MCL1-S, over the antiapoptotic isoform MCL1-L.

The effect of hinokiflavone on MCL1 splicing and its ability to induce cell cycle arrest and/or apoptosis, raise the possibility that either hinokiflavone itself, or a synthetic derivative thereof, could be developed in future as a novel cancer therapeutic. We observed significant variation in the dose response to hinokiflavone between human cancer cell lines. The greatest effect on cell survival was seen in the human acute promyeolytic leukemia cell line NB4, where extensive apoptosis was induced upon treatment with 10 μM hinokiflavone. This positively correlated with the large effect of hinokiflavone in also altering alternative pre-mRNA splicing in NB4 cells, including promoting splicing of the pro-apoptotic isoform of MCL1. However, it is plausible that an increase in PML SUMOylation also contributes to the extreme sensitivity of NB4 cells to hinokiflavone. For example, our MS data in HEK293 cells showed ∼2-fold increased SUMO2 modification of PML after hinokiflavone treatment. It has been shown that the use of arsenic (As_2_O_3_) for therapeutic treatment of acute promyelocytic leukemia (APL) patients is effective because it triggers degradation of PML and oncogenic PML fusion proteins such as PML-RARalpha through promoting their hyper-SUMOylation^38^. We propose that hinokiflavone could have a similar effect through its ability to block SENP activity and therefore stimulate accumulation of hyperSUMOylated proteins.

In summary, this study has identified the plant derived biflavone hinokiflavone as an inhibitor of SUMO protease activity that affects spliceosome assembly and splice site selection, leading to major changes in alternative splicing patterns in human cancer cell lines. Our development of an efficient route for the chemical synthesis of hinokiflavone (King et. al., manuscript in preparation), has allowed us to confirm that the hinokiflavone molecule is the active compound and will facilitate the future investigation of its detailed mechanism of action and exploration of its development as a potential cancer therapeutic. This should help to clarify the relation between SUMO modification and alternative splicing and allow a better understanding of how hinokiflavone exerts such specific effects on increasing SUMOylation of U2 snRNP proteins.

## Experimental Procedures

### Compounds

Hinokiflavone, amentoflavone, cupressuflavone and sciadopitysin were purchased from Extrasynthese (Genay Cedex, France) and isoginkgetin was purchased from MerckMillipore (Darmstadt, Germany).

### Cell culture, RNA isolation and RT-PCR

HeLa, HEK293 and C33A cells were purchased from ATCC and cultured in Dulbecco’s modified Eagle’s supplemented with 10% fetal bovine serum, 2mM glutamine (Life Technologies, Carlsbad, CA, USA) and 100μg/ml streptomycin (100X stock, Life Technologies). Total RNA was extracted from cells using the NucleoSpin RNA II Kit (Macherey-Nagel, Düren, Germany), according to the manufacturer’s instructions. 200 ng of total RNA was reverse transcribed and amplified using the One Step RT-PCR kit (QIAGEN, Hilden, Germany), according to the manufacturer’s instructions. Primer sequences are listed in table 1.

**Table 1.**
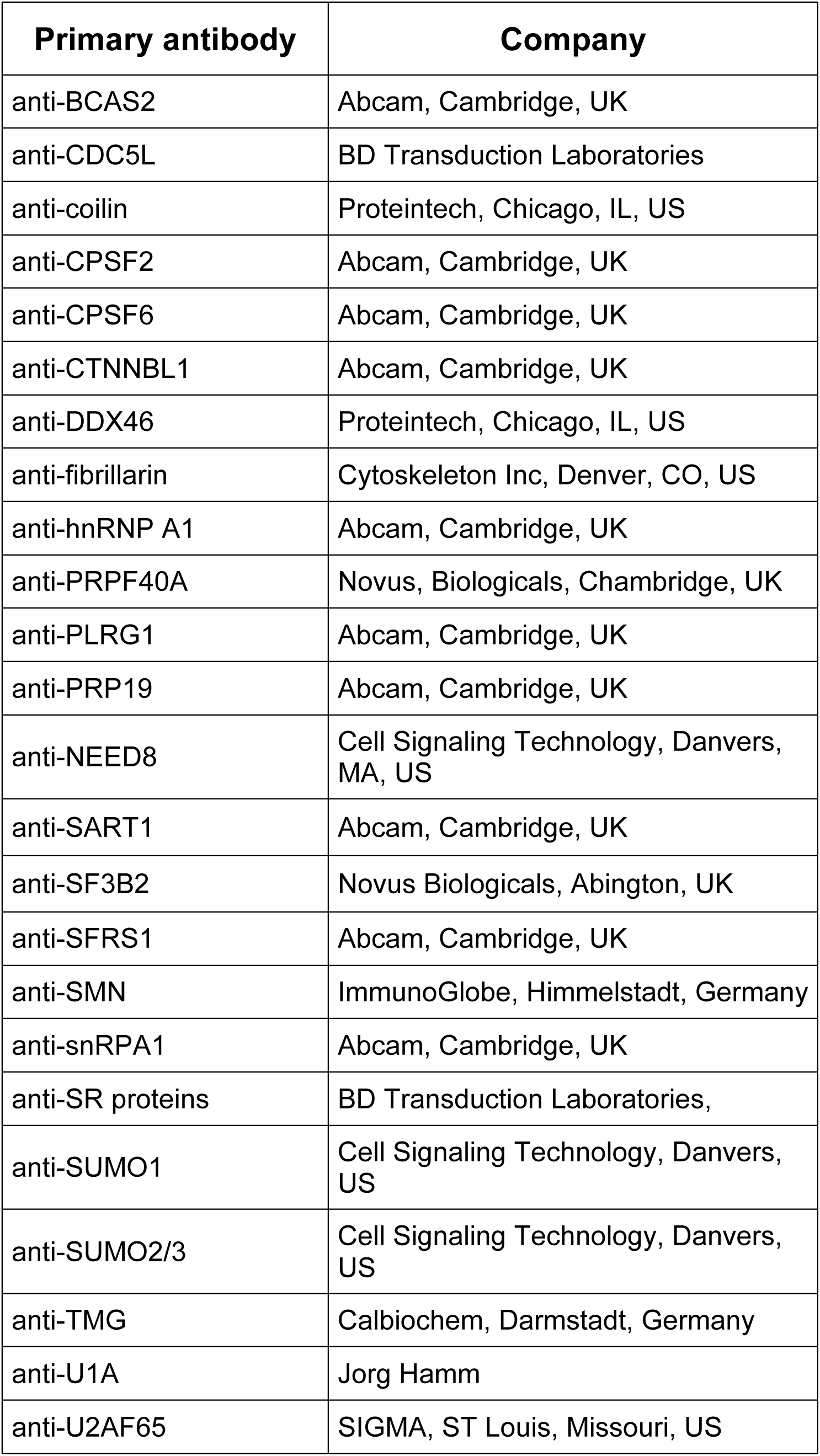
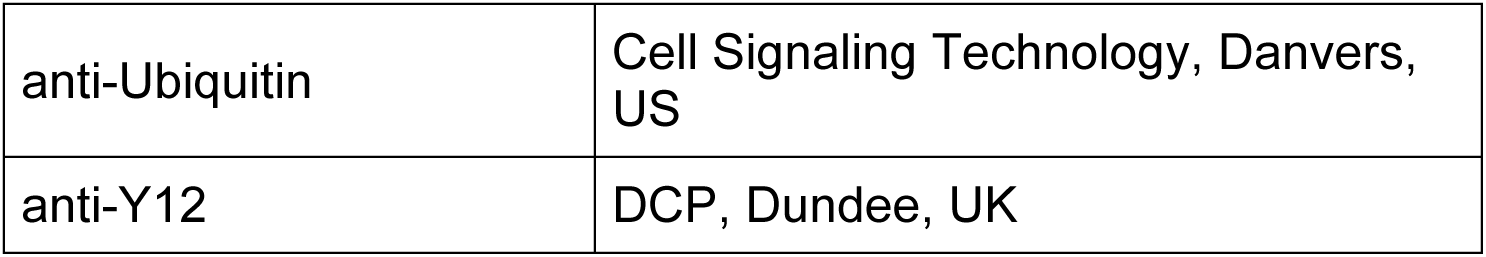

### *In vitro* transcription and RT-based *in vitro* splicing reaction

The Adenovirus pre-mRNA Ad1 and HPV18 E6 pre-mRNA were *in vitro* transcribed using the RNAMAxx High Yield Transcription Kit (Agilent, Santa Clara, CA, USA), according to the manufacturer’s instructions, followed by a DNAase 1 digestion and RNA clean up using the RNeasy Kit (QIAGEN, Hilden, Germany).

Standard splicing reactions were carried out in 30% HeLa nuclear extract (Computer Cell Culture Centre, Seneffe, Belgium), in the presence of either DMSO, or compound, and incubated at 30°C for 90 min. The splicing reaction was followed by a heat inactivation step of 5 min at 95°C before the samples were subjected to proteinase K (QIAGEN, Hilden, Germany) digestion for 30 min at 55°C and another heat inactivation step at 95°C for 5 min. The spliced and non-spliced RNA was amplified using the One step RT-PCR Kit (QIAGEN, Hilden, Germany), according to the manufacturer’s instructions. PCR products were separated by electrophoresis using 1% agarose gels containing SYBR safe DNA gel stain (Life Technologies, Carlsbad, CA, USA).

### Radioactive *in vitro* splicing reaction and native gels

Radioactive *in vitro* splicing reactions were either performed as previously described^1^, using either a ^32^P labelled pBsAd1 pre-mRNA substrate, or a ^32^P labelled MINX pre-mRNA substrate^31^.

Splicing complexes were analysed on a low melting point agarose gel (1.5%, w/v), as previously described^32^ and visualized by phosphor imaging (Typhoon 8600, GE Healthcare, Pittsburgh, PA, USA).

### Gel-based SENP activity assay

SENP activity assays were carried out as 20μl reactions containing 2μl 10x reaction buffer (200 mM Hepes, 500 mM NaCl, 30mM MgCl_2_, pH 7.5), recombinant, *E.coli* expressed SENP1 fragment comprising amino acids 415-643 (186 nM) in SENP buffer (50 mM Tris-HCl, 150 mM NaCl) and 5 μM SUMOylated template YFP-RANGAP (aa 418-587)-ECFP-SUMO2 (5 μM). Either 1 μL DMSO alone (control), or 1 μL compound dissolved in DMSO was added and the reactions were incubated at 37°C for 15 min then stopped by adding 5 μL 4x LDS loading buffer. After incubating the samples at 70°C for 10 min the proteins were separated on a 4-12% Tris-Bis PAGE gel and visualized with Coomassie blue.

### DARTS Assay

To recombinant *E.coli* expressed SENP1 fragment comprising amino acids 415-643 (186 nM) in 20 μl SENP buffer (50 mM Tris-HCl, 150 mM NaCl), either 1 μL DMSO (control), or 1 μL compound dissolved in DMSO, was added and the reactions were incubated at 4°C for 1h. The samples were than treated with different concentrations of pronase (Sigma-Aldrich, St. Louise, MO, US) for 30 min at RT. The reactions were stopped by adding 5 μL 4x LDS loading buffer. After incubating the samples at 70°C for 10 min the proteins were separated on a 4-12% Tris-Bis PAGE gel and visualized with Coomassie blue.

### Cell fixation and Immunofluorescence Analysis

HeLa, HEK293 and NB4 cells were each treated either with DMSO alone (control), or with compound dissolved in DMSO, then grown on cover slips in DMEM for either 4 h, or 24 h, at 37°C before fixing with 4% paraformaldehyde in PHEM buffer for 10 min at RT. After rinsing the cells with PBS, the cells were permeabilized with 0.5% Triton X100 in PBS prior to incubation with the primary antibodies (see table 2). After incubation with the primary antibody for 1 h at RT, the cover slips were washed twice with 0.5% Tween-20 in PBS for 5 min before they were incubated with the dye-conjugated secondary antibody for 30 min. Cells were then stained with DAPI (Sigma-Aldrich, St. Louis, MO, USA) and the cover slips were mounted in Vectashield medium (Vector Laboratories, Peterborough, UK). The samples were visualized using a fluorescence microscope (Zeiss, Jena, Germany; Axiovert-DeltaVision Image Restoration; Applied Precision, LLC).

**Table 2.**
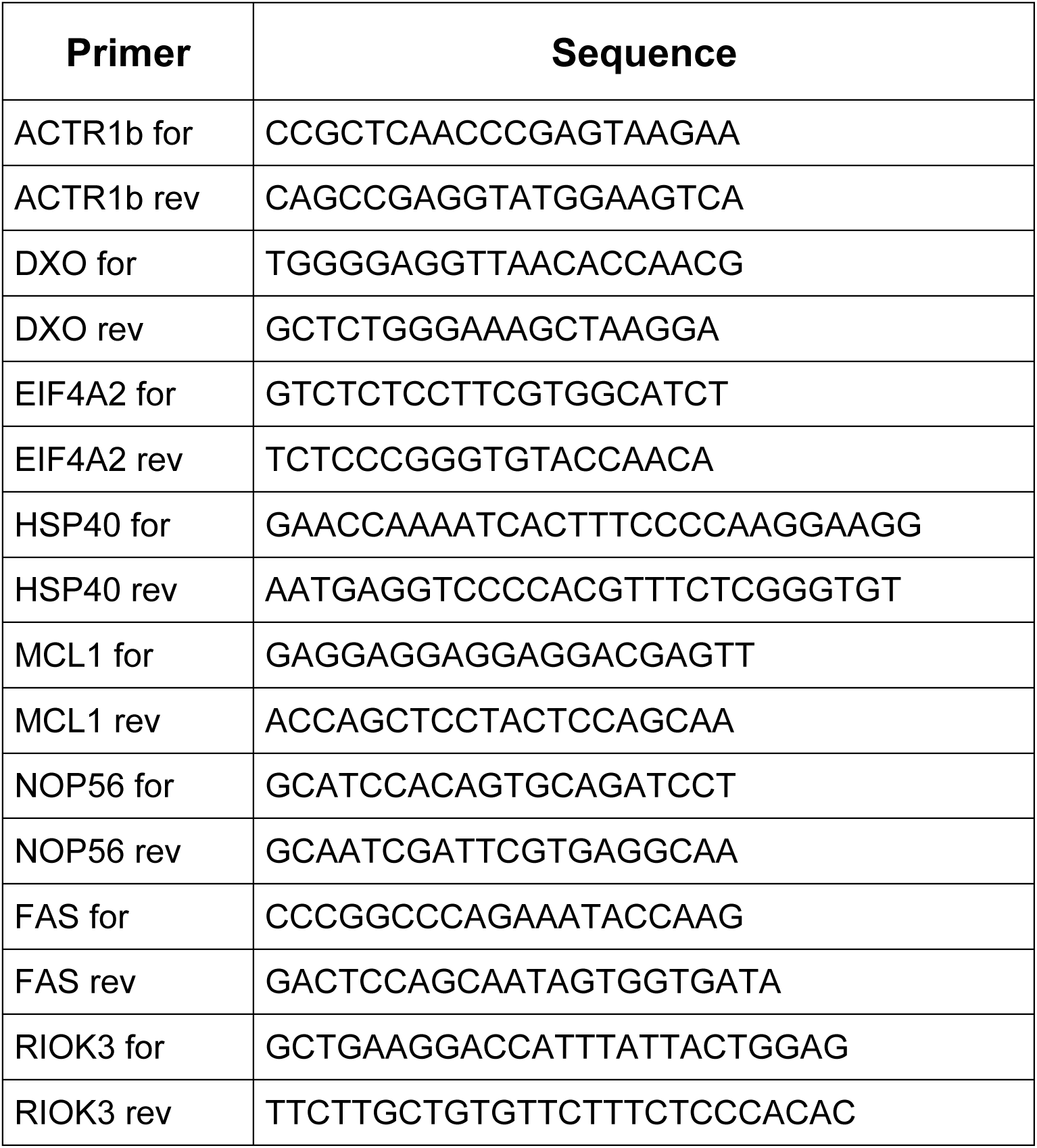

**Table 3.**
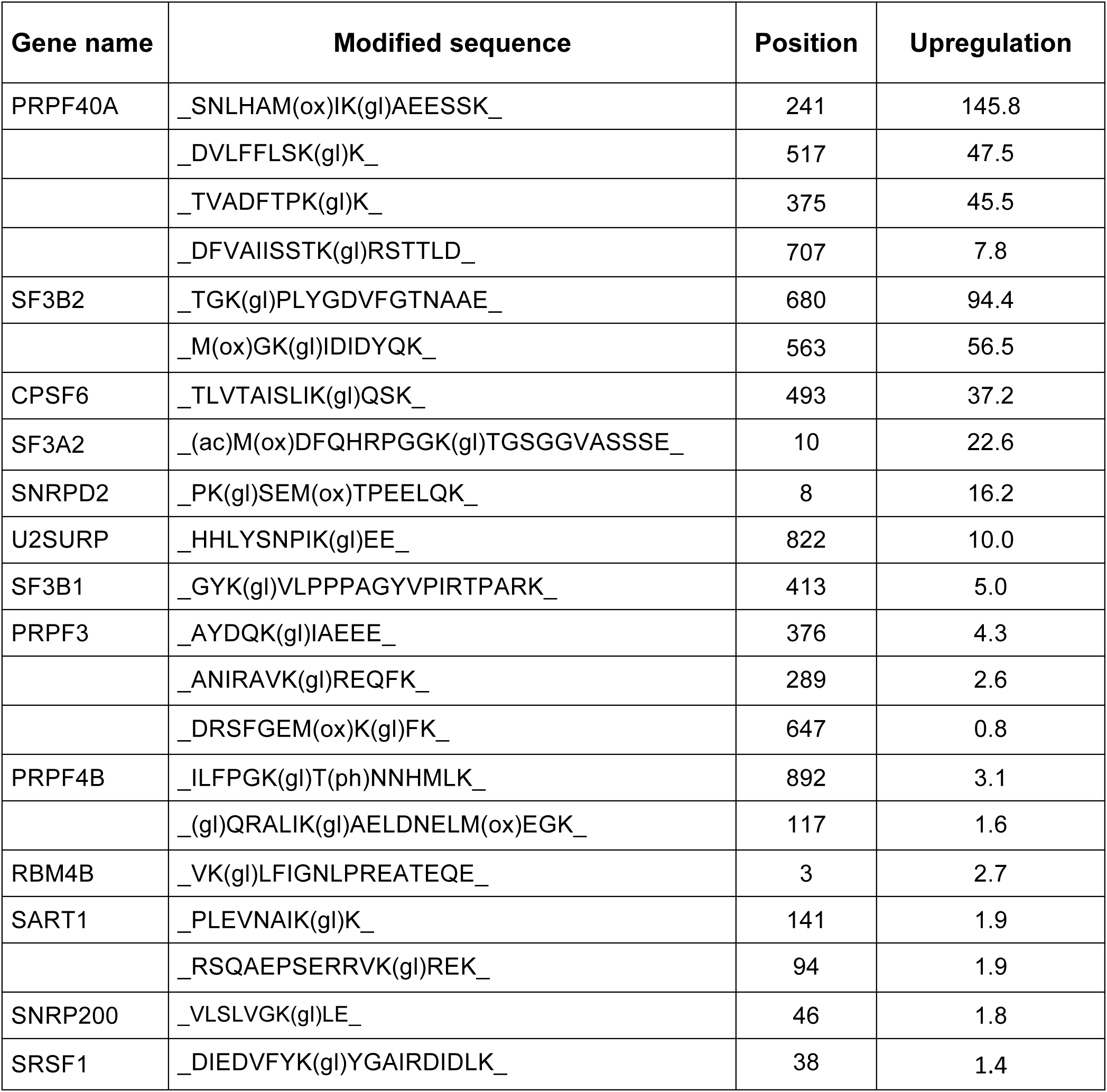

### Flow cytometry analysis

Cells were seeded in 12 well plates and after 24 h were treated with either DMSO alone (control), or compound dissolved in DMSO. The cells were grown at 37°C then harvested after either 4 h, 8 h, or 24 h, then washed twice with PBS, resuspended in cold 70% ethanol and fixed at room temperature for 30 min. Fixed cells were then washed twice with PBS and resuspended in PI stain solution (50 μg/ml propidium iodide and 100 μg/ml ribonuclease A in PBS). Cells were incubated in PI stain solution for 30 min and then analyzed by flow cytometry on a BD FACScalibur. The flow cytometry data were analyzed using FlowJo (TreeStar Inc, Ashland, OR, USA).

### Pulse labeling of HeLa cells with EU

Newly synthesized RNA was detected by using the Click-iT RNA imaging Kit (Life Technologies, Carlsbad, US). In brief, HeLa cells were grown at 37°C for 24h in the presence of either DMSO alone (control), or compound dissolved in DMSO, then pulse labeled for 20 min with EU. The cells were then fixed and the Click-iT detection was performed, according to the manufacturer’s instructions.

### Western blot analysis

Cells treated as described were harvested, washed with PBS and lysed in 1x LDS buffer (Life Technologies, Carlsbad, US). Proteins were separated using a 4-12% NuPAGE Bis-Tris gel (Life, Technologies, Carlsbad, US) and transferred to a nitrocellulose blotting membrane (Amersham, Little Chalfont, UK) by electroblotting. Target proteins were detected with the help of the WesternBreeze^®^ Chemiluminescent Kit, according to the manufacturer’s instructions.

### Cell culture conditions and protein extraction

For quantitative SUMO modification site-specific proteomic experiments, the adherent HEK293^6His-SUMO2-T90K^ N3S cells were cultured in DMEM lacking L-lysine, L-arginine and L-glutamine (Biosera, Uckfield, UK) and supplemented with 10 % dialysed FBS, 1 μg/ml puromycin, 100 U/ml Pen-Strep, 2 mM L-glutamine, and either natural (Sigma-Aldrich, St. Louise, MO, US) (^12^C_6_, ^14^N_2_; K0) and (^12^C_6_, ^14^N_4_; R0), or heavy stable isotope-containing (Cambridge Isotope Laboratories, Tweksbury, US) (^13^C_6_, ^15^N_2_; K8) and CNLM-539-H (^13^C_6_, ^15^N_4_; R10)] 0.8 mM L-lysine and 0.14 mM L-arginine. Five 175 cm^2^ dishes of either light-, or heavy-labelled cells at ∼90 % confluency were transferred into SMEM lacking L-lysine, L-arginine and L-glutamine (Biosera, Uckfield, UK) and supplemented with 10 % dialysed FBS, 1 μg/ml puromycin, 100 U/ml Pen-Strep, 2 mM L-glutamine, and either natural, or heavy stable isotope-containing 0.8 mM L-lysine and 0.14 mM L-arginine, respectively. Approximately 1.0 L of suspension culture, corresponding to ∼4.70 × 10^8^ cells, was cultured per experimental condition and heavy-labelled cells were treated with 20 μM hinokiflavone in DMSO for 8 hours, whereas 20 μM DMSO alone was added to the light-labelled control culture. Cells were harvested by centrifugation, washed with cold 1 × DPBS and 0.9 g of each cell pellet was mixed. The combined pellet of cells was lysed in fresh cell lysis buffer containing 6 M guanidine hydrochloride (Gu-HCl), 100 mM sodium phosphate buffer pH 8.0, 10 mM Tris-HCl pH 8.0, 10 mM imidazole and 5 mM 2- mercaptoethanol. Lysis buffer was added in a ratio of 25:1 (vol/wt). DNA was disrupted by short pulses of sonication, insoluble material was removed by centrifugation and filtration through sterile Minisart NML syringe filters with 0.2 μm pore size (Sartorius, cat. no. 16534) and protein concentration was estimated using Pierce bicinchoninic acid (BCA) protein assay (Thermo Fisher Scientific, Waltham, US).

### Nickel affinity chromatography and protein digestion

Approximately 98.5 % of cell lysate protein was used for the identification of SUMO modification sites and peptides were prepared essentially as previously described^39^. Cell lysate protein was mixed with pre-equilibrated Ni^2+^-NTA agarose resin (QIAGEN, Hilden, Germany) in a ratio of 200:1 (wt/vol) and the slurry was rotated overnight at 4°C. Beads were collected into an empty spin chromatography column (Bio-Rad,) and washed with five resin volumes of cell lysis buffer, ten resin volumes of wash buffer pH 8.0 (8 M urea, 100 mM sodium phosphate buffer pH 8.0, 10 mM Tris-HCl pH 8.0, 10 mM imidazole, 5 mM 2-mercaptoethanol), ten resin volumes of wash buffer pH 6.3 (8 M urea, 100 mM sodium phosphate buffer pH 6.3, 10 mM Tris-HCl pH 8.0, 10 mM imidazole, 5 mM 2-mercaptoethanol) and ten resin volumes of wash buffer pH 8.0. Conjugates were eluted in three sequential steps, with two resin volumes of elution buffer (200 mM imidazole in wash buffer pH 8.0) and protein concentration was estimated by UV-visible spectrophotometry at 280 nm. Protein digestion was performed on an ultrafiltration spin column with a 30 kDa nominal molecular weight cutoff limit (Sartorius, Epsom, UK). Denatured proteins were concentrated on the device, washed twice with 8 M urea, 100 mM Tris-HCl pH 7.5 and treated with 50 mM 2- chloroacetamide in 8 M urea, 100 mM Tris-HCl pH 7.5 at room temperature for 20 minutes in the dark. Samples were washed once with 8 M urea, 100 mM Tris-HCl pH 7.5, twice with IAP buffer (50 mM MOPS-NaOH pH 7.2, 10 mM Na_2_HPO_4_, 50 mM NaCl) and digested overnight with Lysyl endopeptidase (Lys-C; Wako, cat. no. 129-02541) in IAP buffer at 37 °C [enzyme-to-protein ratio 1:50 (wt/wt)]. Peptides were collected and the device was washed with IAP buffer to increase the the yield of Lys-C digested peptides. High-molecular-weight peptides retained on the 30 kDa ultrafiltration spin column were subsequently digested with endoproteinase Glu-C (Sigma-Aldrich, cat. no. 11047817001) overnight in IAP buffer at 25°C [enzyme-to-protein ratio 1:100 (wt/wt)] and after the collection of peptides, the device was washed with IAP buffer.

### Immunoaffinity purification of diGly-Lys-containing peptides

Peptides in IAP buffer were supplemented with ∼18.75 μg of Κ-ε-GG-specific antibody BS3 cross-linked to 3 μl of protein A agarose resin and rotated overnight at 4 °C. Agarose beads were washed twice with 50 resin volumes of cold IAP buffer and peptides were eluted in two sequential steps with 50 μl of 0.15 % (vol/vol) trifluoroacetic acid (TFA).

### In solution digestion of cell lysate proteins

Complete proteome analysis was performed with ∼120 μg of cell lysate protein dissolved in 6 M urea, 2 M thiourea after TCA-based precipitation. The proteins were treated with 10 mM DTT for 1 hour and 50 mM 2-chloroacetamide for 1.5 hours at room temperature in the dark, prior to 2.5-fold dilution into 50 mM ammonium bicarbonate and digestion with Lys-C at enzyme-to-protein ratio 1:50 (wt/wt) at room temperature for 4.5 hours. Lys-C digested samples were then divided in two, and one of the samples was diluted twofold into 50 mM ammonium bicarbonate followed by supplementation with trypsin at enzyme-to-protein ratio 1:50 (wt/wt). Both samples were digested overnight at room temperature. Resulting peptide mixtures were fractionated into six fractions based on the pH of the solution (pH 11.0, 8.0, 6.0, 5.0, 4.0, and 3.0) used to elute the peptides from a pipette tip-based Empore anion exchanger (Agilent Technologies, Santa Clara, US) according to a protocol described in ^40^.

### Liquid chromatography (LC)-tandem mass spectrometry (MSMS)

Prior to MS-based analyses, all peptide samples were desalted on self-made Empore C18 (Agilent Technologies, cat. no. 12145004) Stop and Go Extraction Tips (StageTips) according to a protocol by^41^. Desalted peptide samples were analysed using EASY-nLC 1000 nano-flow UHPLC system, EASY-Spray ion source and Q Exactive hybrid quadrupole-Orbitrap mass spectrometer (all Thermo Fisher Scientific). Peptides were loaded onto 2 cm Acclaim PepMap 100 C18 nanoViper pre-column (75 μm inner diameter; 3 μm particles; 100 Å pore size) at a constant pressure of 800 bar and separated using 50 cm EASY-Spray PepMap RSLC C18 analytical column (75 μm inner diameter; 2 μm particles; 100 Å pore size) maintained at 45°C.

DiGly-Lys-enriched samples were analysed at least twice. Exploratory analysis using standard MS settings was performed using 10 % of the sample and peptides were separated with 60- minute linear gradient of 5-22 % (vol/vol) acetonitrile in 0.1 % (vol/vol) formic acid at a flow rate of 250 nl/min, followed by a 12- minute linear increase of acetonitrile to 40 % (vol/vol). Total length of the gradient including column washout and re-equilibration was 90 minutes. Comprehensive analyses of diGly Lys-containing peptide samples were performed using identical LC conditions. Peptides corresponding to the complete human proteome were analysed with 190-minute linear gradient of 5-22 % (vol/vol) acetonitrile in 0.1 % (vol/vol) formic acid at a flow rate of 250 nl/min, followed by a 25 minute linear increase of acetonitrile to 50 % (vol/vol). Total length of this gradient was 240 minutes.

Peptides eluting from the liquid chromatography column were charged using electrospray ionisation and MS data were acquired online in a profile spectrum data format. Full MS scan covered a mass range of mass-to-charge ratio (m/z) either 300–1800, or 300–1600, during standard and comprehensive peptide analyses, respectively. Target value was set to 1,000,000 ions with a maximum injection time (IT) of 20 ms and full MS was acquired at a mass resolution of 70,000 at m/z 200. Data dependent MS/MS scan was initiated if the intensity of a mass peak reached a minimum of 20,000 ions. During standard LC-MS/MS analyses, up to 10 most abundant ions (Top 10) were selected using 2 Th mass isolation range when centered at the parent ion of interest. For comprehensive analyses, up to 4 (Top 4) most abundant ions were picked for MS/MS. Selection of molecules with peptide-like isotopic distribution was preferred. Target value for MS/MS scan was set to 500,000 ions with a maximum IT of 60 ms and resolution of 17,500 at either m/z 200 for standard, or maximum IT of 500 ms and a resolution 35,000 at m/z 200 for comprehensive peptide analyses. Precursor ions were fragmented by higher energy collisional dissociation (HCD), using normalised collision energy of 30 and fixed first mass was set to m/z 100. Precursor ions with either undetermined, single, or high (>8), charge state were rejected. Ions triggering a data-dependent MS/MS scan were placed on the dynamic exclusion list for either 40 s (standard analyses), or 60 s (comprehensive analyses) and isotope exclusion was enabled. The sample of Lys-C digested diGly-Lys-containing peptides was analysed twice according to the settings of the comprehensive analysis, however the 1,126 peptide ions identified with the first comprehensive LC-MS/MS analysis were added to the exclusion list.

### Analysis of raw MS files

Raw mass spectrometric data files were processed with MaxQuant software (version 1.3.0.5)^42^ and peak lists were searched with an integrated Andromeda search engine^43^ against an entire human UniprotKB proteome containing canonical and isoform sequences^44^ downloaded in April 2013 and supplemented with the primary sequence of 6His-SUMO2^T90K^. Raw files were divided into parameter groups based on the specificity of the proteolysis applied during sample preparation. Hydrolysis of peptide bonds C-terminal to either Lys, or Lys and Arg, with a maximum of three missed cleavages was allowed for peptides processed with either Lys-C, or with Lys-C and Trypsin, respectively. Samples acquired after an additional Glu-C digestion were analysed with enzyme specificity set to C-terminal to Lys, Glu and Asp with a maximum of five missed cleavages. Carbamidomethylation of cysteine residues was specified as a fixed modification and oxidation of methionines, acetylation of protein N-termini and where applicable, Gly-Gly adduct on internal lysine residues, were selected as variable modifications. Additional analyses were performed by including either deamidation of Gln and Asn, or conversion of N-terminal Glu or Gln to pyroglutamate as extra variable modifications. Maximum peptide mass of 10,000 Da was allowed, multiplicity was set to 2, and K8 and R10 were selected as heavy-labelled counterparts. A maximum of either six, or four, labelled residues were allowed per peptide during the analyses of either diGly-Lys-enriched samples, or peptides representing the entire proteome, respectively. A decoy sequence database was generated using Lys as a special amino acid when analysing diGly-Lys-containing peptides. Default values were chosen for the rest of the parameters. All datasets were filtered by posterior error probability to achieve a false discovery rate of 1 % at protein, peptide and modified site level. The non-normalised SILAC ratios of diGly-Lys-containing peptides were normalised according to the median normalisation factor obtained for the complete proteome experiment and was based on the non-normalised and normalised protein ratios reported in the MaxQuant output file.

### Stable cell lines

cDNA of PRPF40A was purchased from GenScript (Piscataway, US) and cloned into the pEGFP-C3 vector using EcoRI/BamHI restriction sites. HEK293 cell line stably expressing GFP-PRPF40A were generated by transfecting the constructs using Effectene (QIAGEN, Hilden, Germany), according to the manufacturer’s instructions and maintaining the cells in DMEM (low glucose), with 10% FBS and 0.5 mg/ml G418 (Invitrogen, Carlsbad, USA).

### Co-Immunoprecipitation

HEK293 cells stably expressing GFP-PRPF40A were seeded into 10 cm dishes for 24h before the cells were treated either with DMSO alone (control), or 20 μΜ hinokiflavone in DMSO, for 8h before cells were trypsinised and harvested. Cells were washed twice with ice-cold PBS before ∼ 1×10^7^ cells were lysed in 1 ml of Co-IP buffer (1mM EDTA, 100μΜ Na_3_VO_4_, 0.5% Triton X-100, 20mM beta-Glycerol P), with protease inhibitors and NBE for 30 min. Before adding 50μl of GFP-Trap magnetic beads (Chromotek, Apple Vally, US) to 1 ml of cleared lysate, the lysed cells were centrifuged for 10 min at 13.0000rpm at 4°C. After incubation at 4°C overnight, the beads were washed 3 times with Co-IP lysis buffer and proteins were eluted from the beads by adding 200μl 1×LDS buffer and boiling the samples for 10 min at 95°C. The samples were subjected to western blot analysis with the indicated primary antibodies.

## Acknowledgements

We thank our colleagues for advice and discussion. We thank staff in the Fingerprints, Flow Cytometry and Cell Sorting Facility for advice and assistance (CAST, University of Dundee). This work was supported by a Wellcome Trust programme grant to AIL (073980/Z/03/B) and also supported by infrastructure funded by a Wellcome Trust Strategic award (097045/B/11/Z). R.T.H. is a Senior Investigator of the Wellcome Trust (grant 098391/Z/12/7) T.T. was funded by the EU Initial Training Network (ITN) ‘‘UPSTREAM’’ (PITN-GA-2011-290257) and an ISSF grant funded by the Wellcome Trust (105606/Z/14/Z).

## Supplementary Figures

**Supplementary Figure 1. Amentoflavone, Cupressoflavone and Sciadopitysin do not alter pre-mRNA splicing *in cellulo*.** HEK293 cells were treated with either DMSO (control), or with 20 μΜ, 40 μM, 60 μM, 80 μM, or 100 μM of either amentoflavone, cupressoflavone, or sciadopitysin, for 24h. Using RT-PCR and primer pairs to detect changes in exon skipping of RBM5, FAS and MCL1 pre-mRNAs, as well as intron inclusion in HSP40 and RIOK3 pre-mRNAs, no alteration of pre-mRNA splicing for these transcripts were detected in the presence of these bioflavonoids.

**Supplementary Figure 2. Fas pre-mRNA splicing in the presence of hinokiflavone**. Schematic representation of Fas isoforms formed in hinokiflavone treated cells.

**Supplementary Figure 3. Comparison of natural and synthetic hinokiflavone**. Proton nuclear magnetic resonance spectrum of natural (red) and synthetic (blue) hinokiflavone (A). Effect of natural and synthetic hinokiflavone on the alternative splicing of the MCL1 gene in HEK293 cells (B).

**Supplementary Figure 4. Hinokiflavone treatment leads to an increase in SUMO2/3 modified proteins in HeLa and NB4 cells**. HeLa and NB4 cells were treated with either DMSO (control), or with 0.5 μM, 1 μM, 2.5 μM, 5 μM, 10 μM, 20 μM, or 30 μM of hinokiflavone for 24h. SUMO2/3 and alpha tubulin expression were detected by western blotting.

